# Seeded assembly *in vitro* does not replicate the structures of α-synuclein filaments from multiple system atrophy

**DOI:** 10.1101/2020.11.27.401364

**Authors:** Sofia Lövestam, Manuel Schweighauser, Shigeo Murayama, Yuko Saito, Taisuke Tomita, Takashi Ando, Kazuko Hasegawa, Mari Yoshida, Airi Tarutani, Masato Hasegawa, Michel Goedert, Sjors H.W. Scheres

## Abstract

The propagation of conformational strains by templated seeding is central to the prion concept. Seeded assembly of α-synuclein into filaments is believed to underlie the prion-like spreading of protein inclusions in a number of human neurodegenerative diseases, including Parkinson's disease, dementia with Lewy bodies (DLB) and multiple system atrophy (MSA). We previously determined the atomic structures of α-synuclein filaments from the putamen of five individuals with MSA. Here, we used filament preparations from three of these brains for the *in vitro* seeded assembly of recombinant human α-synuclein. We find that the structures of the seeded assemblies differ from those of the seeds, suggesting that additional, as yet unknown, factors play a role in the propagation of pathology. Identification of these factors will be essential for understanding the prion-like spreading of α-synuclein proteinopathies.

## Introduction

The ordered assembly of a small number of proteins into pathological amyloid filaments defines most neurodegenerative diseases, including Alzheimer disease (AD) and Parkinson disease (PD) (Goedert, 2015). Diseases characterised by the assembly of α-synuclein and tau are the most common proteinopathies of the human nervous system. Most cases of disease are sporadic, but a small percentage is inherited.

The first assemblies form in a small number of cells in a given brain region, from where they spread through prion-like mechanisms (Goedert, 2015). A central tenet of the prion hypothesis is that proteinopathies are characterised by assemblies with specific conformations that propagate from cell to cell (Goedert et al., 2010; Prusiner, 1982). Spreading is consistent with staging schemes that have postulated a stereotypical progression of inclusions from single sites (H. Braak & Braak, 1991; Heiko Braak et al., 2003). Decades elapse between the formation of assemblies and the appearance of disease symptoms, providing an important therapeutic window. Evidence for the existence of prion-like mechanisms in human brain has come from the development of scattered α-synuclein inclusions in foetal human midbrain neurons that were therapeutically implanted into the striata of patients with advanced PD (Kordower et al., 2008; J.-Y. Li et al., 2008).

α-Synuclein assemblies are characteristic of PD, PD dementia, DLB, MSA, and several rarer conditions, known collectively as synucleinopathies (Goedert et al., 2017). In these diseases, the 140 amino acid α-synuclein assembles into a filamentous, β-sheet-rich conformation. Unbranched α-synuclein filaments are 5-10 nanometres in diameter and up to several micrometres in length. They are found mostly in nerve cells (Lewy bodies and neurites) and, for MSA, also in glial cells, chiefly in oligodendrocytes (glial cytoplasmic inclusions, GCIs, or Papp-Lantos bodies). Filamentous α-synuclein is phosphorylated and exhibits additional posttranslational modifications (Fujiwara et al., 2002; Sorrentino & Giasson, 2020), but it remains to be shown that these modifications are necessary for assembly. Amino acids 30-100 have been reported to make up the structured part of α-synuclein filaments (Miake et al., 2002). A seed of α-synuclein can trigger the assembly of soluble α-synuclein (Luk et al., 2009; Yonetani et al., 2009).

A link between α-synuclein assembly and disease was established by the findings that missense mutations in SNCA (the α-synuclein gene), and multiplications of this gene, cause rare forms of inherited PD and PD dementia (Polymeropoulos et al., 1997; Singleton et al., 2003). Some SNCA mutations and gene multiplications also cause DLB. Abundant α-synuclein inclusions are present in all cases of inherited disease. Sequence variation in the regulatory region of SNCA is associated with increased α-synuclein expression and a heightened risk of developing sporadic PD, which accounts for over 90% of cases of this disease (Nalls et al., 2014). Expressed α-synuclein, wild-type or mutant, assembles into filaments *in vitro* (Conway et al., 1998). Moreover, expression of human mutant α-synuclein in animal models causes its aggregation and neurodegeneration (Giasson et al., 2002).

Experimental evidence has shown that assembled α-synuclein from MSA behaves like a prion (Holec & Woerman, 2020). Intracerebral or peripheral injection of MSA brain extracts into heterozygous mice transgenic for human A53T α-synuclein led to the formation of abundant neuronal α-synuclein inclusions and their spreading, accompanied by motor impairment (Lavenir et al., 2019; Watts et al., 2013; Woerman et al., 2015, 2018). Protein misfolding cyclic amplification (PMCA) and real time-induced quaking induced conversion (RT-QuIC), have been reported to discriminate between MSA and PD (Shahnawaz et al., 2020).

Following the identification of α-synuclein filaments from DLB by negative-stain immuno-electron microscopy (immuno-EM) (Spillantini et al., 1998), multiple techniques, including solid-state nuclear magnetic resonance, electron diffraction, X-ray diffraction and electron cryo-microscopy (cryo-EM), have been used to study the molecular structures of recombinant α-synuclein filaments (Guerrero-Ferreira et al., 2018, 2019; Rodriguez et al., 2015; Serpell et al., 2000; Shahwanaz et al., 2020; Strohäker et al., 2019; Tuttle et al., 2016; Vilar et al., 2008). In some of these studies, filaments were also amplified by using seeds from human brain and recombinant human protein as substrate.

We recently showed that the structures of α-synuclein filaments from MSA consist of type I and type II filaments, each with two different protofilaments (Schweighauser et al., 2020). By two-dimensional class averaging, filaments from the brains of individuals with MSA differ from those of DLB, suggesting that distinct strains do indeed characterise synuclein proteinopathies. However, as is the case of tau assemblies (Scheres et al., 2020), the structures of α-synuclein filaments from brain are unlike those formed from recombinant proteins. The main differences are in the extended folds of MSA protofilaments, their asymmetrical packing and the presence of non-proteinaceous molecules between protofilaments.

These findings raised the question if seeded assemblies of α-synuclein have the same structures as those of brain seeds. Here, we show that the cryo-EM structures of seeded recombinant α-synuclein assemblies differ from those of MSA seeds. This suggests that in disease, additional molecules and/or posttranslational modifications of α-synuclein are required for the faithful replication of filament structures.

## Results

### Seeded assembly of α-synuclein with filament preparations from MSA brains

We seeded the *in vitro* assembly of recombinant wild-type human α-synuclein with filament preparations from the putamen of three cases of MSA (Materials & Methods). The cryo-EM structures of the filaments from these cases are known (cases 1, 2 and 5 in Schweighauser et al., 2020). They contain variable proportions of type I and type II MSA filaments, with I:II ratios of 80:20 for case 1; 20:80 for case 2; and 0:100 for case 5. We monitored the kinetics of aggregation using thioflavin T (Xue et al., 2017). The assembly conditions were as described (Shahnawaz et al., 2020), using 100 mM piperazine-N,N′-bis(2-ethanesulfonic acid) (PIPES) and 500 mM NaCl at 37° C, pH 6.5. Upon addition of seeds, we observed a lag phase of 20 - 40 hrs, before fluorescence increased rapidly and plateaued after 30 – 60 hrs (Figure 1b). Case 5 seeds were faster at seeding recombinant α-synuclein and resulted in higher fluorescence intensities than seeds from cases 1 and 2 No increase in fluorescence was observed in the absence of seeds. Negative-stain EM confirmed the presence of abundant filaments after incubation with MSA seeds (Figure 1c).

**Figure 1.**
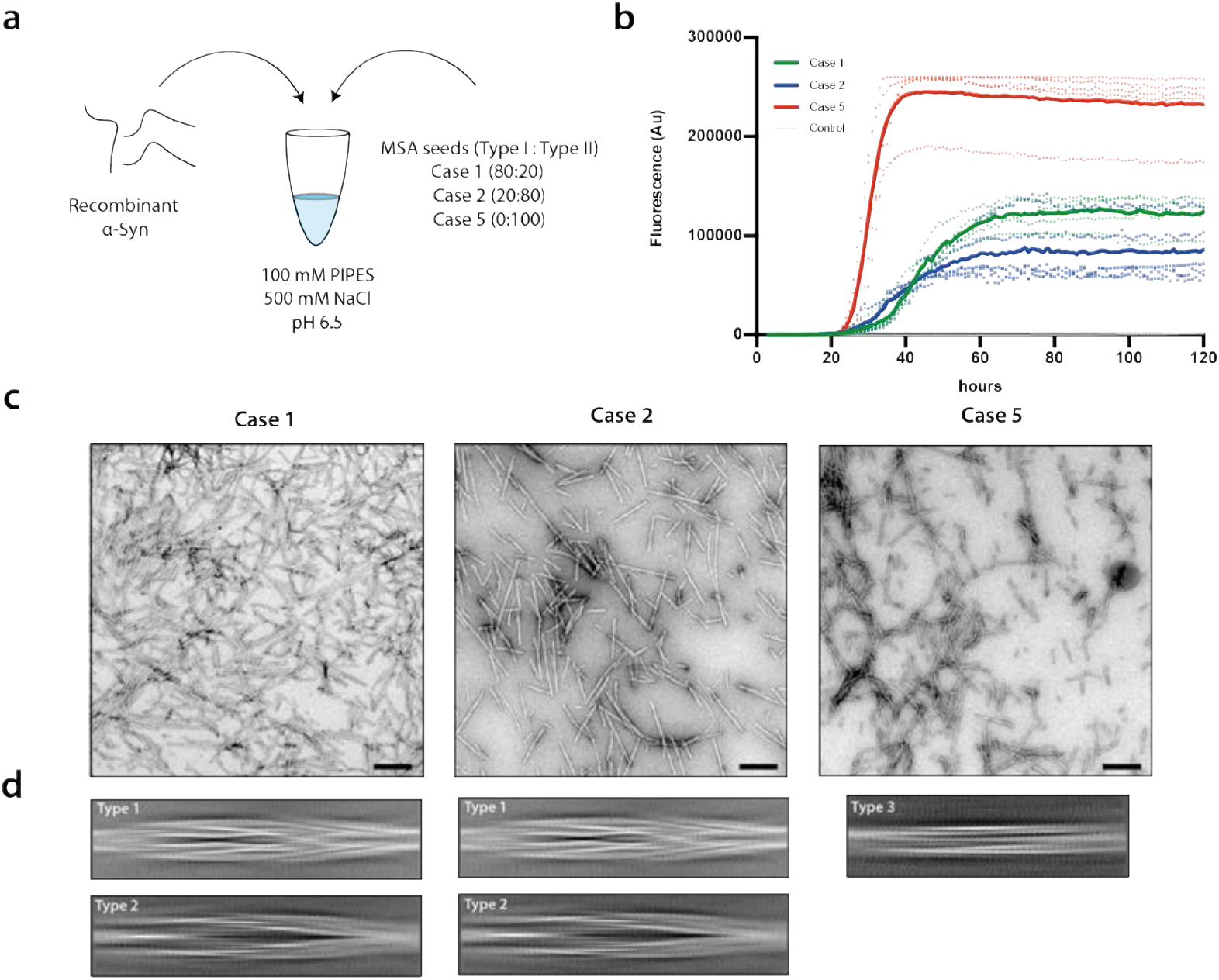
Seeded assembly of recombinant α-synuclein with filament preparations from MSA brains. **(a)**Recombinant wild-type human α-synuclein was mixed with sonicated MSA seeds in 100 mM PIPES, 500 mM NaCl, 0.05% NaN3, pH 6.5. Seeds had variable ratios of type I and type II filaments. **(b)**Assembly was quantitated by thioflavin T fluorescence of recombinant α-synuclein in the presence of MSA seeds from case 1 (green), case 2 (blue) and case 5 (red). Controls (grey) were without seeds. Curves represent the mean and dots correspond to the values in each experiment, (n=5). **(c)** Negative stain micrographs of α-synuclein filaments after seeded assembly. **(d)** Cryo-EM 2D class averages in boxes spanning 825 Å of the types of filaments. Assembly with seeds from MSA cases 1 and 2 gave rise to type 1 and type 2 filaments. Type 3 filaments formed when the seeds were from MSA case 5.

### Cryo-EM imaging of seeded α-synuclein filaments

We used cryo-EM to image the filaments formed following incubation of recombinant α-synuclein with seeds from each MSA case. Visual inspection of micrographs of filaments from experiments that used seeds from MSA cases 1 and 2 indicated the presence of two main filament types, which we called type 1 and type 2. Type 1 filaments have an average crossover distance of 800 Å and widths of 60-130 Å; type 2 filaments have a crossover distance of 900 Å and widths of 80-130 Å. We also observed straight filaments with no observable twist. It is unclear if they correspond to filaments of types 1 or 2 that untwisted because of sample preparation artefacts, such as interactions with the air-water interface, or if they represent additional filament types. Due to the lack of twist, we were unable to solve the structures of these filaments.

Two-dimensional classification readily separated type 1 and type 2 filaments for further processing and indicated that both types are 2-fold symmetric along their helical axis (Fig. 1d). Further 3D classification revealed that type 1 and type 2 filaments occurred in two variants in the data set of filaments that formed with seeds from MSA case 1. They are characterised by small differences in protofilament folds. We called the predominant protofilament ‘fold A’ and the minor protofilament ‘fold B’. We could not identify protofilaments with fold B when seeds from MSA case 2 were used. Using helical reconstruction in RELION (He & Scheres, 2017), we determined cryo-EM structures of type 1 and type 2 filaments with only protofilament fold A to 3.4 Å resolution (Figure 2; Figure 2 - figure supplements 1 and 2). Reconstructions of type 1 and type 2 filaments with two protofilaments of fold B, or with one protofilament of fold A and another protofilament of fold B, were solved to resolutions of 3.4 – 4.1 Å (Figure 3; Figure 3 - figure supplement 1). Reconstructions of filaments containing protofilaments of fold B were less well defined than those of filaments with two protofilaments of fold A. Assembly with seeds from MSA case 5 resulted almost exclusively in the formation of a different type of filament, which we called type 3. Type 3 filaments were thinner, more bendy and longer than filaments of types 1 and 2. Type 3 filaments have a crossover of 900 Å and widths of 55-65 Å. We solved their structure to 3.2 Å resolution (Figure 4; Figure 4 - figure supplements 1 and 2). A minority of filaments (< 2%) comprised a doublet of the type 3 filaments. Throughout this manuscript, we use blue colours for fold A and green for fold B of type 1 and type 2 filaments, and we use purple for type 3 filaments.

**Figure 2.**
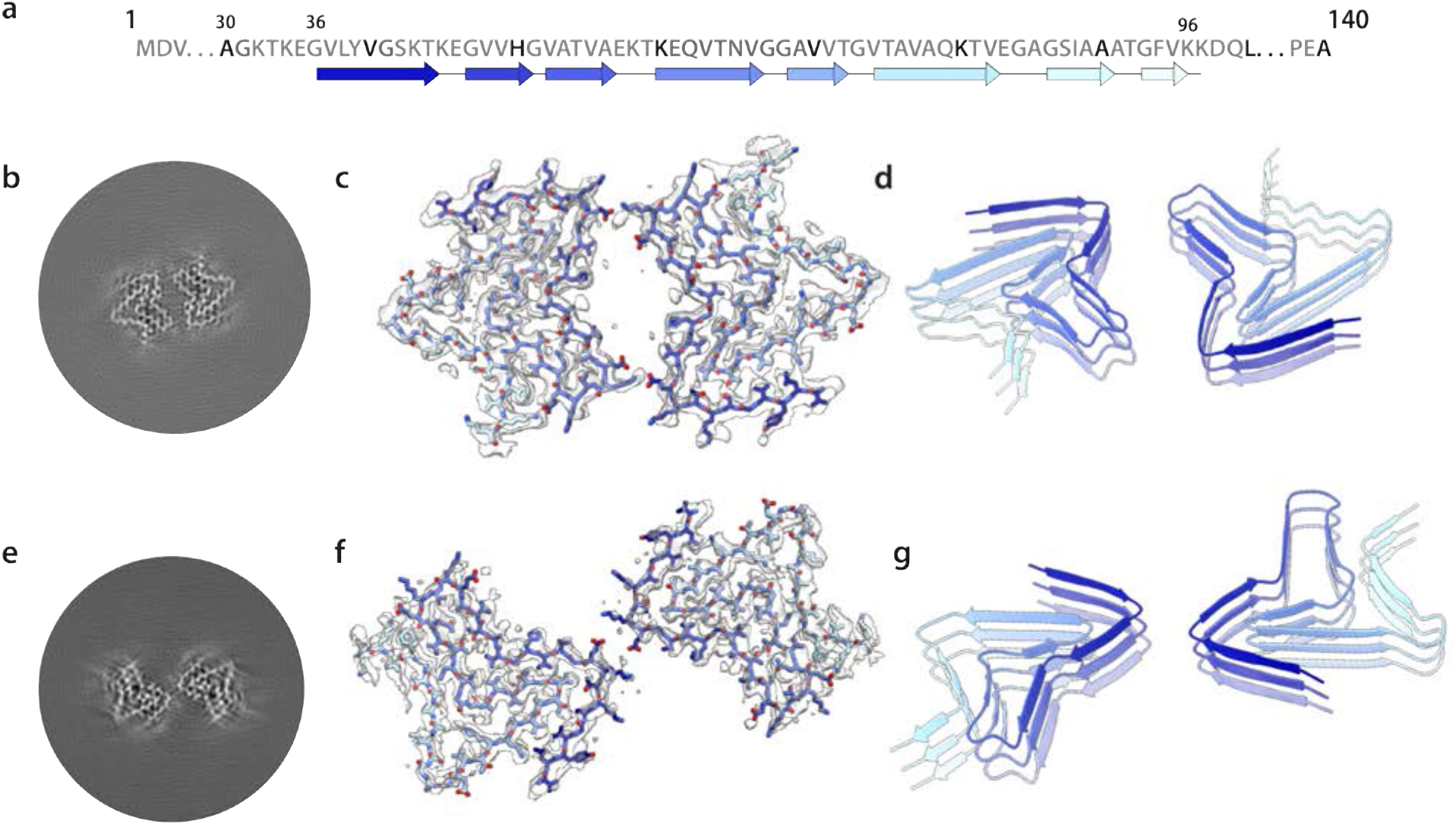
Cryo-EM structures of type 1 and type 2 filaments with protofilament fold A assembled using seeds from MSA case 2. **(a)** Primary sequence of α-synuclein with β-strands and loop regions shown from dark blue (N-terminal) to light blue (C-terminal). **(b)** Central slice of the 3D map for type 1 filaments with protofilament fold A. **(c)** Cryo-EM density (transparent grey) and fitted atomic model (with the same colour scheme as in a) for type 1 filaments. **(d)** Cartoon view of three successive rungs of the type 1 filament. (e-g) As (b-d), but for type 2 filaments.

**Figure 3.**
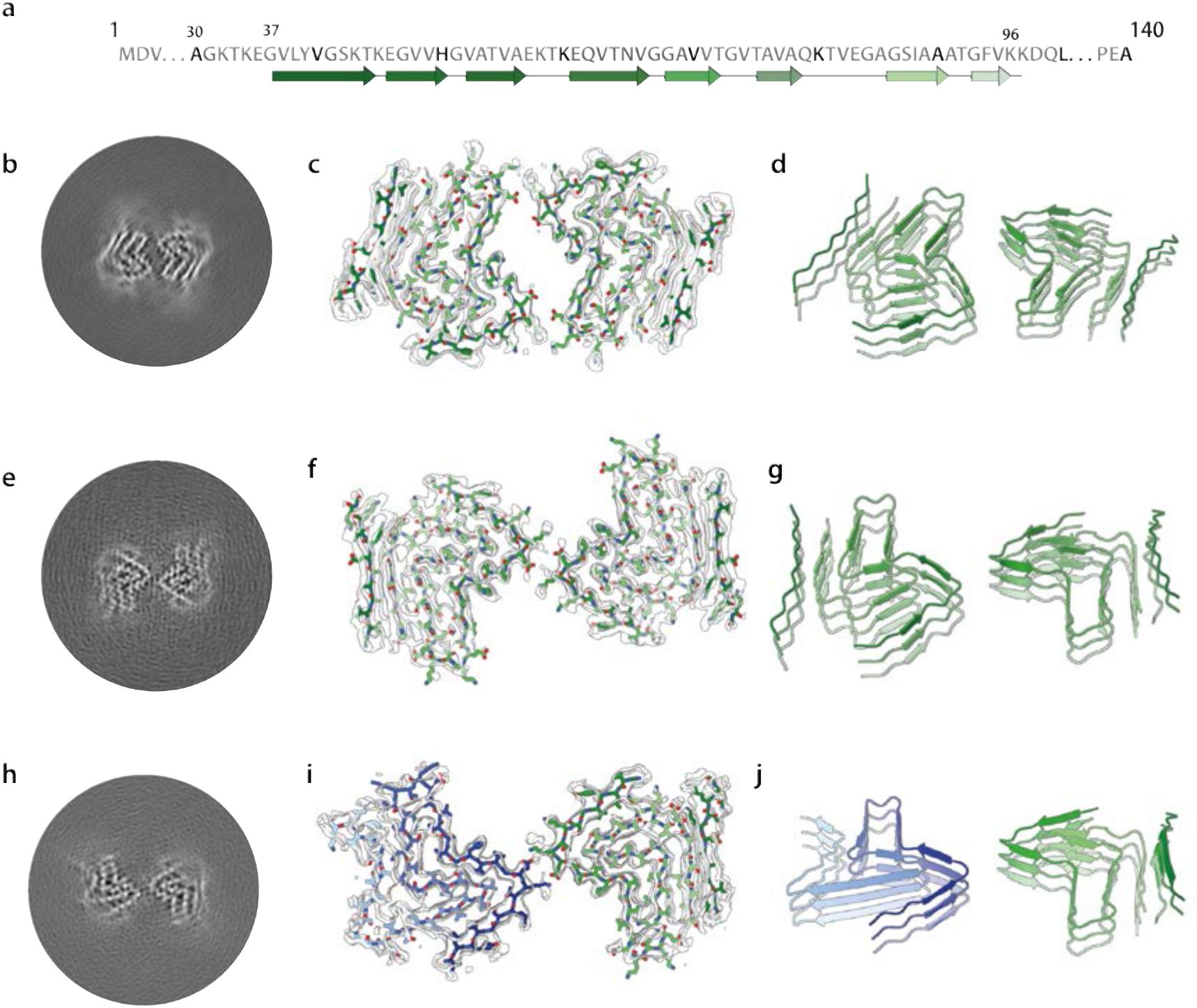
Cryo-EM structures of of type 1 and type 2 filaments with protofilament fold B assembled using seeds from MSA case 1. Central slice through the reconstruction of the type 1 filament with protofilament fold B (left) and an overlay of the density (in transparent grey) and the atomic model (right). **(b)** As in (a), but for the type 2 filament. **(c)** As in (a), but for the putative type 2 filament that contains a mixture of protofilament folds A and B.

**Figure 4.**
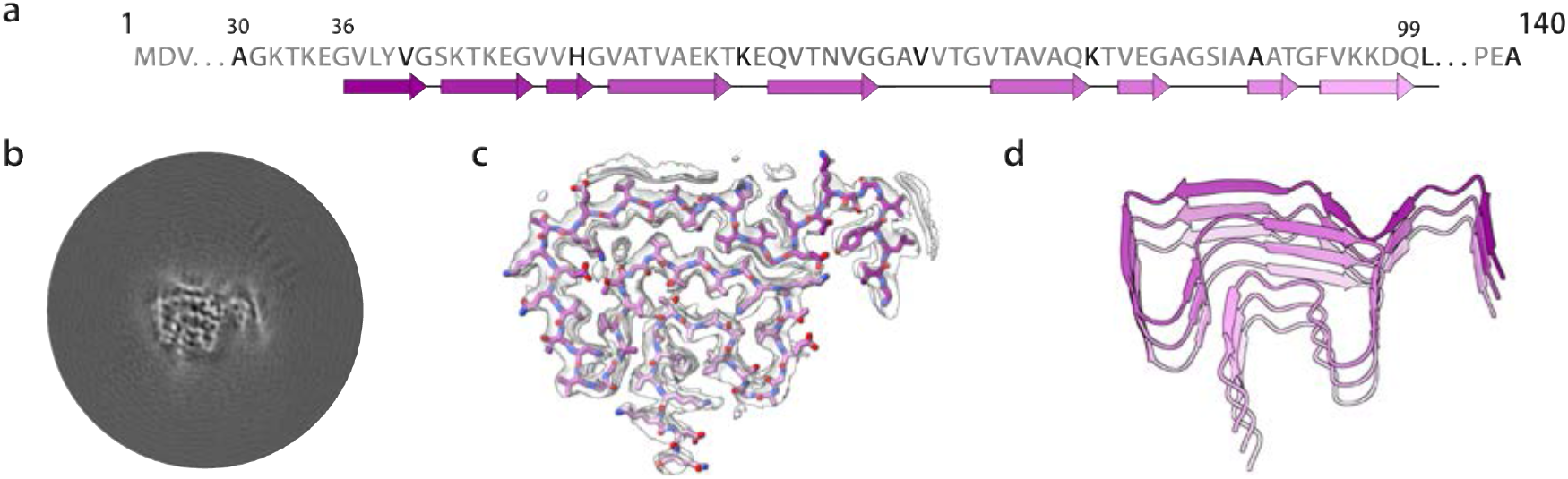
Cryo-EM structure of type 3 filaments assembled using seeds from MSA case 5. **(a)** Primary sequence of α-synuclein with β-strands and loop regions shown from dark violet (N-terminal) to light pink (C-terminal). **(b)** Central slice of the 3D map for the type 3 filament. **(c)** Cryo-EM density (transparent grey) and the fitted atomic model (with the same colour scheme as in a). **(d)** Cartoon view of three successive rungs of the type 3 filament.

### Cryo-EM structures of type 1 and type 2 α-synuclein filaments

Most type 1 and type 2 filaments that formed with seeds from MSA case 1, and all the filaments that formed with seeds from MSA case 2, consisted of two protofilaments of fold A that were related by C2 symmetry. Filaments of types 1 and 2 differed in their inter-protofilament packing (Figure 2). In type 1 filaments, two salt bridges between E46 and K58 held the protofilaments together, by creating a large solvent-filled channel. The inter-protofilament interface in type 2 filaments was formed by two salt bridges between K45 and E46 of each protofilament. The smeared reconstructed densities at the points furthest away from the helical axis suggest that the inter-protofilament interface of type 2 filaments is more flexible than that of type 1 filaments. Protofilament fold A consists of 8 β-sheets: β1-6 form a roughly Z-shaped hairpin-like structure, with glycines or KTK motifs between the β-sheets at the bends; β7-8 fold back against β4, leaving a small triangular cavity between β5, β6 and β7. This fold is unlike any of those of the MSA type I and type II protofilaments. It is almost identical to the protofilament fold that was reported for *in vitro* aggregated recombinant E46K α-synuclein (Boyer et al., 2020), although the inter-protofilament interface was different from the interfaces observed here for type 1 and type 2 filaments (Figure 2 - figure supplement 3). A minority of type 1 and 2 filaments that formed with seeds from MSA case 1 consisted of two symmetry-related copies of protofilaments with fold B. Although the reconstructions of type 1 and type 2 filaments with two protofilaments of fold B (Figure 3; Figure 3 - figure supplement 1) were less well defined than those for filaments with two protofilaments of fold A, the maps revealed that fold B is nearly identical with the structure of filaments assembled from wild-type recombinant α-synuclein (Guerrero-Ferreira et al., 2019). This increased our confidence in building and refining an atomic model for the protofilaments with fold B. The resulting model from the type 2 filament has a root-mean-square-deviation (r.m.s.d.) of 1.38 Å with the structure of assembled wild-type α-synuclein (Guerrero-Ferreira et al., 2019). Again, protofilament fold B was unlike any of the four protofilaments from MSA type I and type II filaments. An asymmetric reconstruction from a subset of the images suggested that asymmetric type 2 filaments may also form from one protofilament with fold A and one protofilament with fold B (Figure 4c). However, we cannot exclude the possibility that this reconstruction is an artefact arising from suboptimal classification of filament segments. Folds A and B are almost identical at residues G36–V55, and V63–A78, with some flexibility in the β-turn at residues E57-E61. However, comparing the more compact fold B to fold A, a flip in K80 from the hydrophobic core towards the solvent results in a sharp turn at T81 and a shift by three residues in the packing of β4 against β7 (Figure 3 - figure supplement 3).

### Cryo-EM structure of type 3 α-synuclein filaments

Type 3 filaments consist of a single protofilament that extends from G36-Q99 and comprises 10 β-sheets (β1-10) (Figure 4). Residues 46-99 form a Greek key motif, as described before (Tuttle et al., 2016),with a salt bridge between E46 and K80. This motif is preceded by a β-arch formed by residues Y39-T44 and Y39-E46. The density between residues 36 and 39 is more smeared. Two stretches of elongated, smeared densities, possibly originating from parts of the N-terminus of α-synuclein, are observed in front of β1 in the β-arch and β4 in the Greek key motif. An additional fuzzy density is observed in front of the side chains of K43, K45 and H50. Whereas filament types 1 and 2 did not resemble the four protofilaments observed in MSA, type 3 filaments were almost identical to protofilament IIB2, with an r.m.s.d. between atomic coordinates of 1.02 Å (Figure 5). However, in MSA filaments, K58 is flipped away from the core of the protofilament to form a salt bridge with T33 of the opposing protofilament, whereas K58 forms part of the protofilament core in type 3 filaments. Minor rearrangements occur near V40, which is also involved in inter-protofilament packing in MSA filaments. Interestingly, the position of the density of the unidentified co-factor at the inter-protofilament interface of type II filaments coincides with the fuzzy density in front of K43, K45 and H50. Type 3 filaments are almost identical to the narrow protofilament formed upon *in vitro* assembly of recombinant H50Q α-synuclein (Boyer et al., 2019), with an r.m.s.d. between atomic coordinates of 0.62 Å (Figure 5 - figure supplement 1).

**Figure 5.**
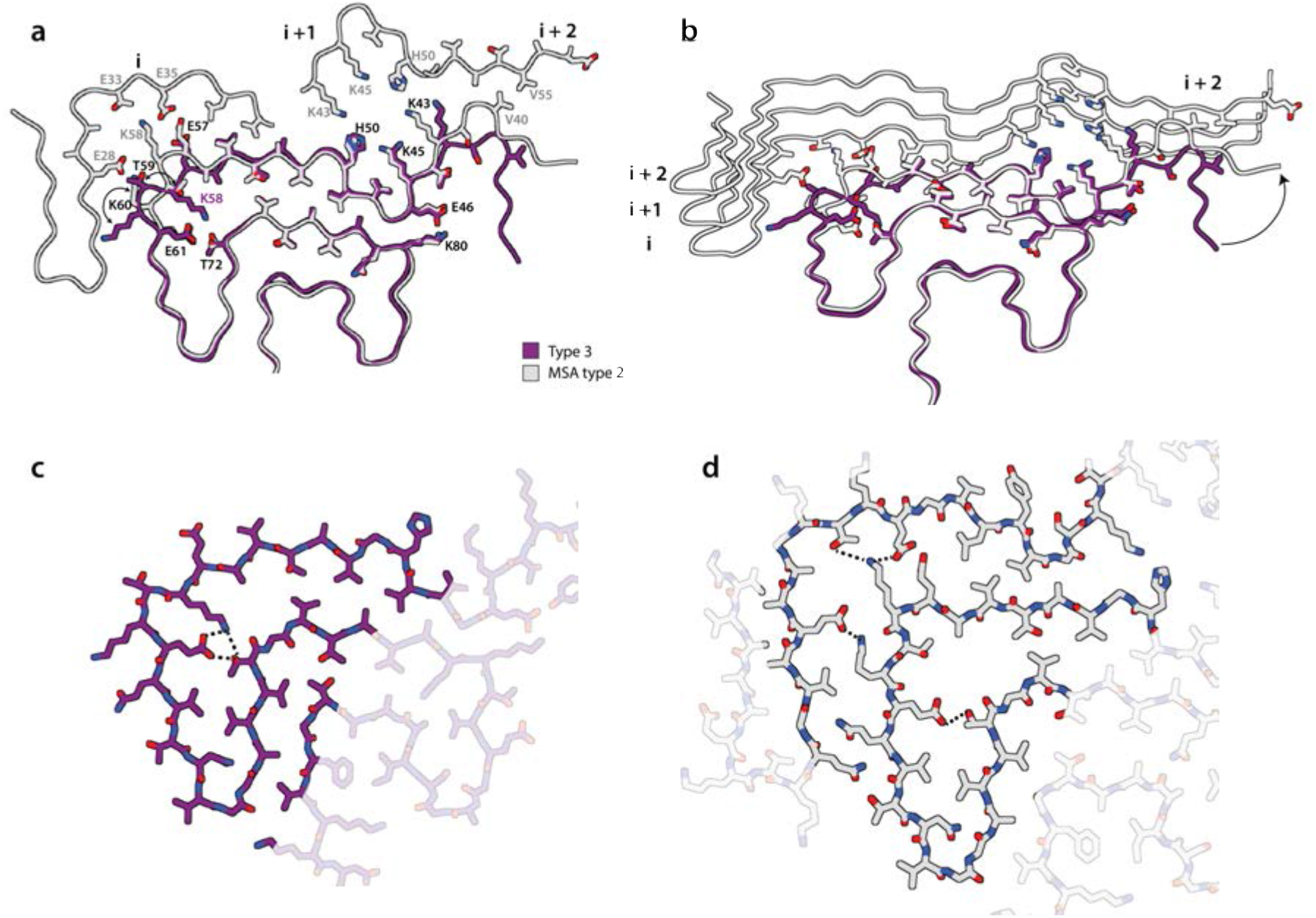
Comparison of type 3 filament with protofilament IIB from MSA case 5. Atomic model of the type 3 filament (purple) overlaid with the model of protofilament IIB2 from MSA case 5. The additional density at the protofilament interface of MSA type II filaments is shown in orange. (b) Cartoon view of one rung of type 3 filaments overlaid with one rung of protofilament IIB and three rungs of protofilament IIA of MSA case 5. Residues on MSA protofilament IIA that interact with the rung of protofilament IIB shown are highlighted with sticks. **(c)** Close up all-atom view of the hydrogen-bonding network (yellow dashed) between K58, E61 and T72 in type 3 filaments. (d) As in (c), but for protofilaments IIA and IIB in MSA filaments.

### Cryo-EM structures of α-synuclein filaments from second-generation seeded aggregation

To further explore the effects of buffer conditions on seeded aggregation, we incubated seeds from MSA case 5 with recombinant human α-synuclein in phosphate-buffered saline (PBS). We previously observed that the density for the additional molecules at the interface between protofilaments in our reconstructions of MSA filaments (Schweighauser et al., 2020) overlaps with similar densities in reconstructions of *in vitro* aggregated recombinant α-synuclein, which have been attributed to phosphate ions (Guerrero-Ferreira et al., 2018, 2019). Since the additional density in MSA filaments could accommodate two phosphate ions, we supplemented PBS with 1 mM pyrophosphate. However, by negative-stain imaging, the seeded assemblies were indistinguishable from those formed using PBS without pyrophosphate. We then performed second-generation seeded assembly, in which the aggregates from the assembly in PBS-pyrophosphate were used as seed. Cryo-EM structure determination of the seeded assemblies confirmed the faithful propagation of type 3 filaments, with a larger proportion of type 3 doublet filaments (~5%) (Figure 4 - figure supplement 2).

## Discussion

We show here that the structures of seeded assemblies of wild-type recombinant human α-synuclein differ from those of seeds that were extracted from the brains of individuals with MSA (Figure 6). We used the assembly conditions of Shahnawaz et al. (2020) who reported that PMCA, using cerebrospinal fluid as seed and recombinant α-synuclein as substrate, can discriminate between PD and MSA. It remains to be seen if α-synuclein seeds from PD brain yield structures that are different from those described here. Nevertheless, our results raise important questions for the study of amyloid structures and prion processes.

**Figure 6.**
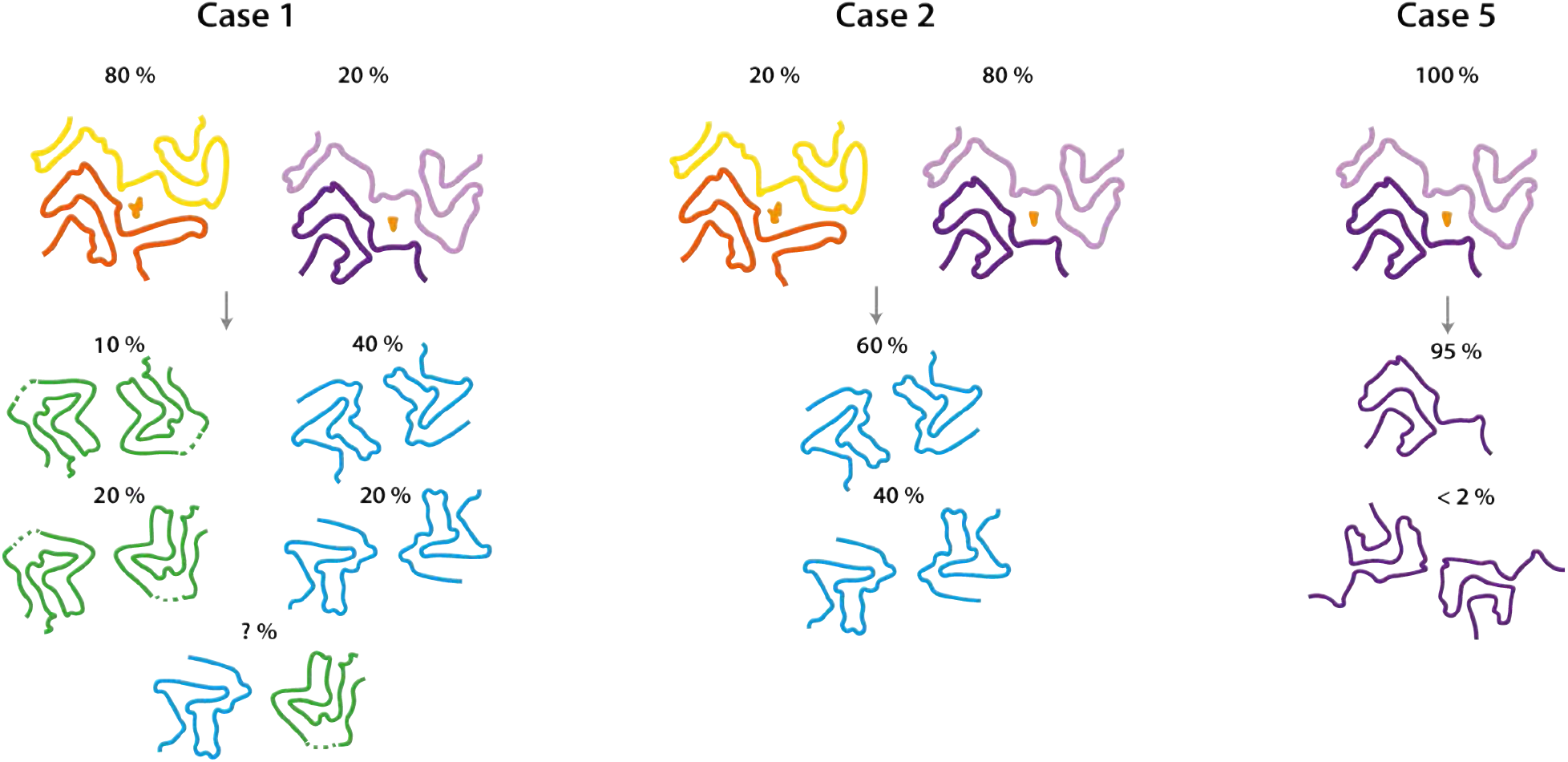
Summary of MSA seeded aggregation experiments. Cartoon illustrations show the structures of MSA type I and type II filaments and their relative quantities in MSA cases 1, 2 and 5 at the top, and the products of seeded aggregation underneath.

Amyloid filaments are structurally versatile, with the same amino acid sequences being able to adopt different structures (Guerrero-Ferreira et al., 2020; Scheres et al., 2020). Moreover, the cryo-EM structures of tau, β-amyloid and α-synuclein filaments from human brain are different from those of recombinant proteins assembled *in vitro* (Fitzpatrick et al., 2017; Kollmer et al., 2019; Schweighauser et al., 2020). The present findings demonstrate that, even when using brain-derived filament preparations to seed *in vitro* assembly, the resulting structures are unlike those of the seeds.

When using seeds from MSA cases 1 and 2, which contain a mixture of type I and type II filaments, and recombinant human α-synuclein as substrate, we observed the formation of type 1 and type 2 filaments. When using seeds from MSA case 5, with only type II filaments, we observed the formation of filaments of type 3. These observations suggest that in seeded assemblies, type I filaments overshadow type II MSA filaments, despite the observation that seeds of case 5 resulted in a faster and stronger increase in thioflavin-T fluorescence compared to seeds from cases 1 and 2. The possibility that different conformational strains have different seeding potencies has implications for the interpretation of prion propagation assays.

It is commonly assumed that self-propagation of strains occurs through templated incorporation of monomers at the ends of amyloid filaments. Indeed, following sonication, α-synuclein filaments had increased seeding potencies (Tarutani et al., 2016, 2018). However, it is unclear how this could explain the formation of type 1 and type 2 filaments with markedly different protofilament folds, when compared to MSA filaments. Each prion strain is believed to comprise a large number of conformationally distinct assemblies (also known as clouds), often with a dominant conformer that propagates under host selection (Collinge & Clarke, 2007; J. Li et al., 2010). Our work on tau and α-synuclein assemblies has shown the presence of only one or two major filament types in the brains from patients at end-stage disease Scheres et al., 2020; Schweighauser et al., 2020). It is possible that type 1 and type 2 filaments were present in the filament preparations from MSA brains, but not numerous enough to be detected by cryo-EM (Schweighauser et al., 2020). We previously demonstrated that tau structures that only made up around 3% of filaments can be detected (Falcon et al., 2019), indicating that, if present in MSA brains, type 1 and type 2 α-synuclein filaments are infrequent.

Type 3 filaments, which assembled from MSA type II seeds, fit the model of structural equivalence between seeds and seeded assemblies better than type 1 and type 2 filaments, because their structure overlaps almost completely with that of type IIB protofilaments from the putamen of patients with MSA. We previously attributed additional cryo-EM densities at the inter-protofilament interfaces of type I and type II MSA filaments to negatively charged, non-proteinaceous molecules. It is possible that the absence of these molecules in the seeded assembly experiments led to the formation of a structure that represents only half of the seed structures. These findings indicate that protofilament IIB, but not IIA, can form from recombinant α-synuclein through seeded assembly without added cofactor.

Abundant GCIs in oligodendrocytes are the major neuropathological hallmark of MSA (Papp et al., 1989). Thus, differences in the cellular milieu between oligodendrocytes and other brain cells may play a role in the seeded aggregation of MSA filaments. Oligodendrocytes have been shown to transform misfolded α-synuclein into a GCI-like strain (Peng et al., 2018).

Besides the possible incorporation of other molecules in α-synuclein filaments from human brain, it is also conceivable that recombinant α-synuclein is not able to form MSA filaments. Truncation and post-translational modifications of α-synuclein may be needed (Fujiwara et al., 2002; Sorrentino & Giasson, 2020). In α-synuclein filament preparations from the putamen of patients with MSA, mass spectrometry identified N-terminal acetylation, C-terminal truncation, ubiquitination at K6 K12, K21, acetylation at K21 K23 K32 K34 K45 K58 K60 K80 and K96 and phosphorylation at Y39, T59, T64, T72 and T81 (Schweighauser et al., 2020). It is not known if these modifications occur prior to, during or after filament assembly, and if or how they may affect filament conformations. Assembly of recombinant wild-type human α-synuclein using seeds of α-synuclein phosphorylated at Y39 gave rise to filaments with a different fold from that of the seeds (Zhao et al., 2020). Moreover, C-terminal truncation of recombinant α-synuclein has been shown to promote filament assembly *in vitro* (Crowther et al., 1998); inhibiting C-terminal truncation in transgenic mouse models of MSA has been reported to reduce pathology (Bassil et al., 2016; Sorrentino and Giasson, 2020). It has also been shown that interactions with lipids, DNA, RNA, iron and phosphate promote α-synuclein aggregation *in vitro*, and similar interactions could be important for the formation of MSA filaments in brain (Buell et al., 2014; Galvagnion et al., 2016; Ostrerova-Golts et al., 2000).

Identification of the factors that govern the replication of conformational prion strains will be essential for our understanding of propagation of the distinct proteinopathies. Meanwhile, the relevance of the structures of amyloids assembled from recombinant protein seeds and the results of self-propagation studies should be interpreted with care.

## Acknowledgements

We thank the families of the patients for donating brain tissues; T. Nakane for help with RELION; W. Zhang and Y.Shi for helpful discussions; T. Darling and J. Grimmett for help with high-performance computing. M.G. is an Honorary Professor in the Department of Clinical Neurosciences of the University of Cambridge and an Associate Member of the UK Dementia Research Institute. This work was supported by the UK Medical Research Council (MC-U105184291 to M.G. and MC_UP_A025_1013, to S.H.W.S.), Eli Lilly and Company (to M.G.) and the Japan Agency for Medical Research and Development (JP18ek0109391 and JP18dm020719, to M.H.). This study was supported by the MRC-LMB electron microscopy facility.

## Author contributions

S.L. performed seeded aggregation and cryo-EM experiments and analysed the data, with contributions from M.S., M.G. and S.H.W.S.; Y.S., S.M., T.T.,T.A., K.H., M.Y., A.T. and M.H. identified patients, performed neuropathology and extracted α-synuclein filaments from MSA cases; S.H.W.S. and M.G. supervised the project; S.L., M.G. and S.H.W.S. wrote the manuscript, with inputs from all authors.

## Ethical review processes and informed consent

The procedures for the extraction of MSA filaments from human brain were approved through the ethical review process at Tokyo Metropolitan Institute of Medical Science. Informed consent was obtained from the patients’ next of kin.

## Materials and Methods

### Expression and purification

α-Synuclein was expressed and purified, essentially as described (Morgan et al., 2020). Briefly, plasmid pRK172 encoding a cDNA for full-length, wild-type human α-synuclein was transformed into *E*. *coli* BL21(DE3)-gold (Agilent Technologies). Cells were cultured in 2xTY, 5mM MgCl2 and 100 mg/l ampicillin at 37 ºC until an OD600 of 0.7 was reached; α-synuclein expression was then induced with 1 mM IPTG. After 4 hrs, cells were harvested by centrifugation and resuspended in buffer A [50 mM Tris-HCl, pH 7.5, 10 mM EDTA, 2.5 mM TCEP (Sigma-Aldrich), 0.1 mM AEBSF (Sigma-Aldrich), 40 μg/ml DNase and 10 μg/ml RNase (Sigma-Aldrich), supplemented with cOmplete EDTA-free Protease Inhibitor Cocktail (Roche)]. They were lysed by sonication on ice using a Sonics VCX-750 Vibra Cell Ultra Sonic Processor for 5 min (5 s on, 10 s off) at 40 % amplitude. The lysates were centrifuged at 17,000 x g for 40 min at 4 °C, filtered with a 0.45 μM cut-off filter, loaded onto an anion exchange Sepharose 26/10 Q column (GE Healthcare) and eluted with a 0-1 M NaCl gradient. Fractions containing α-synuclein were precipitated using ammonium sulphate (0.3 g / ml) for 30 min at 4 °C and centrifuged at 16,000 g for 30 min at 4 ° C. The resulting pellets were resuspended in buffer B (PBS, 0.1 mM AEBSF, supplemented with cOmplete EDTA-free Protease Inhibitor Cocktail), loaded onto a HiLoad 16/60 Superdex (GE Healthcare) column equilibrated in buffer B and eluted using a flow rate of 1 ml/min. The purity of α-synuclein was analysed by SDS-PAGE and protein concentrations determined spectrophotometrically using an extinction coefficient of 5600 M^−I^ cm^−I^.

### Extraction of MSA filament seeds

The filament preparations used in this study have been described (Schweighauser et al., 2020). Briefly, frozen putamen from MSA cases 1, 2 and 5 was homogenised in 20 % vol (w/v) extraction buffer (10 mM Tris-HCl, pH 7.5, 0.8 M NaCl, 1 mM EGTA, 10% sucrose, 2% sarkosyl, pH 7.5) and incubated for 30 min at 37 °C. The homogenates were centrifuged for 10 min at 10,000g at room temperature, followed by a 20 min spin of the resulting supernatants at 100,000g. The pellets were resuspended in 500 μl/g extraction buffer and centrifuged at 3,000g for 5 min to remove large contaminants. The supernatants were diluted in 50 mM Tris-HCl, pH 7.5, containing 150 mM NaCl, 10% sucrose and 0.2% sarkosyl, and centrifuged at 166,000g for 30 min. Sarkosyl-insoluble pellets were resuspended in 50 μl/g tissue and filament concentrations estimated by negative-stain EM. Prior to seeded assembly experiments, pellets were centrifuged at 2,000 g for 5 min, the resulting supernatants were diluted 10-fold, and sonicated in an Eppendorf tube using a VialTweeter (Hielscher) at a cumulative power of 100 W. Sonication did not alter the structure of the seeds, as suggested by negative-stain EM (Figure 1 - figure supplement 1), and as confirmed by cryo-EM 2D class averages of the seeds before and after sonication (Figure 1 - figure supplement 2).

### Seeded assembly

Purified recombinant α-synuclein was centrifuged at 20,000 x g for 1 hr to remove potential aggregates. 70 μM recombinant α-synuclein was incubated with 2 μM MSA seeds (as assessed by negative-stain EM) in 100 mM PIPES pH 6.5, 500 mM NaCl, 0.05% NaN3, and 5 μM thioflavin-T, in a final volume of 200 μl per experiment. Controls used buffer without seeds. Seeded assembly proceeded for 120 h at 37 °C in a FLUOstar Omega (BMG Labtech) microplate reader where the samples were alternatingly shaken for 1 minute at 400 rpm, and left to rest for 1 minute, during which fluorescence was measured.

For cryo-EM, seeded assembly conditions were identical, but no thioflavin-T was added to the buffer and the samples were shaken continuously for 72 hrs. Seeded assembly experiments for cryo-EM were also performed in PBS buffer, supplemented with 1 mM pyrophosphate and 0.05% NaN3. The resulting filaments were pelleted, resuspended in 200 μl and sonicated as described above, and then used as seeds (2 μM) for a second-generation seeded assembly experiment with recombinant α-synuclein (70 μM) in the same PBS buffer.

### Cryo-EM grid preparation and imaging

Prior to freeze plunging, filaments were pelleted for 45 min at 100,000x g and resuspended at 100 μM α-synuclein in 50 mM Tris, pH 7.5, 50 mM NaCl. Four μl of sample was applied to glow-discharged 1.2/1.3 holey carbon coated gold grids (Quantifoil AU R1.2/1.3, 300 mesh) for 30s, blotted with filter paper for 3.5 s and plunge-frozen in liquid ethane using an FEI Vitrobot Mark IV. Filaments were imaged on a Thermo Fischer Titan Krios microscope operating at 300 kV equipped with a Gatan K2 Summit direct detector in counting mode and a GIF Quantum energy filter (Gatan) with a slit width of 20 eV to remove inelastically scattered electrons. Acquisition details are given in Tables 1 and 2.

**Table 1.** Cryo-EM data collection, refinement and validation statistics.

| **Data collection and processing** | Type 1A<br>Case 2<br>(EMDB-xxxx)<br>(PDB<br>xxxx) | Type 2A<br>Case 2<br>(EMDB-xxxx)<br>(PDB xxxx) | Type 1B<br>Case 1<br>(EMDB-xxxx)<br>(PDB xxxx) | Type 2B<br>Case 1<br>(EMDB-xxxx)<br>(PDB xxxx) | Type 2A/B<br>Case 1<br>(EMDB-xxxx)<br>(PDB xxxx) | Type 3<br>Case 5<br>(EMDB-xxxx)<br>(PDB xxxx) | Type 3<br>Case 5<br>Second<br>generation | Type 3 doublet<br>Case 5<br>Second<br>generation |
| --- | --- | --- | --- | --- | --- | --- | --- | --- |
| Magnification | X105 000 | X105 000 | X105 000 | X105 000 | X105 000 | X105 000 | X105 000 | X105 000 |
| Voltage (kV) | 300 | 300 | 300 | 300 | 300 | 300 | 300 | 300 |
| Detector | K2 | K2 | K2 | K2 | K2 | K2 | K2 | K2 |
| Electron exposure (e-/Å<sup>2</sup>) | 32.6 | 32.6 | 36.7 | 36.7 | 36.7 | 37.5 | 37.0 | 37.0 |
| Defocus range (µm) | -1.5 to -2.8 | -1.5 to -2.8 | -1.5 to -2.8 | -1.5 to -2.8 | -1.5 to -2.8 | -1.5 to -2.8 | -1.5 to -2.8 | -1.5 to -2.8 |
| Pixel size (Å) | 1.14 | 1.14 | 1.1 | 1.1 | 1.1 | 1.14 | 1.14 | 1.14 |
| Micrographs | 1294 | 1294 | 2172 | 2172 | 2172 | 1265 | 1317 | 1317 |
| Symmetry imposed | C2 | C2 | C2 | C2 | C1 | C1 | C1 | C2 |
| Initial particle images (no.) | 287 364 | 287 364 | 441 592 | 441 592 | 441 592 | 122 831 | 270 003 | 270 003 |
| Final particle images (no.) | 67 619 | 82 474 | 33 479 | 87092 | 57 358 | 69 490 | 18 691 | 82 474 |
| Map resolution (FSC=0.143) (Å) | 3.47 | 3.43 | 3.84 | 3.55 | 4.23 | 3.18 | 3.54 | 4.40 |
| Map resolution range (Å) | 2.8 – 11 | 3.2 – 6.3 | 3.5 - 10 | 3.3 – 18 | 4.0 - 14 | 2.7 – 5.5 | NA | NA |
| Helical twist (°) | -1.04 | -0.95 | -0.86 | -0.77 | -0.86 | -0.95 | -0.95 | -1.52 |
| Helical rise (Å) | 4.75 | 4.75 | 4.78 | 4.75 | 4.80 | 4.75 | 4.75 | 4.75 |
| **Refinement** |  |  |  |  |  |  |  |  |
| Initial model used (PDB code) | 6UFR | 6UFR | 6SSX | 6SST | 6SST/6UFR | 6PEO |  |  |
| Model resolution (FSC=0.5)(Å) | 3.4 | 3.7 | 5.4 | 4.6 | 5.4 | 3.5 |  |  |
| Map sharpening *B* factor (Å<sup>2</sup>) | -79.5 | -68.3 | -105.7 | -81.9 | -107.8 | -56.6 |  |  |
| Model composition |  |  |  |  |  |  |  |  |
| Non-hydrogen atoms | 5052 | 5032 | 5496 | 5496 | 5274 | 2652 |  |  |
| Protein residues | 732 | 732 | 816 | 816 | 774 | 384 |  |  |
| Ligands | 0 | 0 | 0 | 0 | 0 | 0 |  |  |
| R.m.s. deviations |  |  |  |  |  |  |  |  |
| Bond lengths (Å) | 0.011 | 0.012 | 0.013 | 0.011 | 0.009 | 0.010 |  |  |
| Bond angles (°) | 1.966 | 2.133 | 1.606 | 2.118 | 1.432 | 2.002 |  |  |
| Validation |  |  |  |  |  |  |  |  |
| MolProbity score | 0.88 | 1.12 | 1.03 | 1.12 | 1.06 | 0.97 |  |  |
| Clashscore | 0.00 | 0.38 | 0.00 | 0.27 | 0.18 | 0.18 |  |  |
| Poor rotamers (%) | 0.19 | 0.00 | 0.19 | 0.00 | 0.76 | 0.37 |  |  |
| Ramachandran plot |  |  |  |  |  |  |  |  |
| Favored (%) | 94.49 | 92.23 | 90.62 | 91.15 | 92.01 | 94.09 |  |  |
| Allowed (%) | 5.51 | 7.77 | 9.38 | 8.85 | 7.72 | 5.91 |  |  |
| Disallowed (%) | 0.00 | 0.00 | 0.00 | 0.00 | 0.27 | 0.00 |  |  |

### Helical reconstruction

Filaments were reconstructed in RELION-3.1 (Zivanov et al., 2020) using helical reconstruction (He & Scheres, 2017). Movie frames were corrected for beam-induced motions and dose-weighted in RELION using its own motion-correction implementation (Zivanov et al., 2018). Non-dose-weighted micrographs were used for CTF estimation with CTFFIND-4.1 (Rohou & Grigorieff, 2015). Filaments were picked manually, ignoring those without a clear twist. Initially, particle segments were extracted using a box size of 550 pixels and an interbox distance of 14 Å and downscaled to 225 pixels for 2D classification. For filaments formed from the seeds of MSA cases 1 and 2, filament types 1 and 2 were separated at this initial 2D classification stage. Crossover-distances were obtained by manual measurements in the micrographs and used to calculate initial estimates for the helical twist of the different filament types: −1.0° for type 1; −0.8° for type 2; and −1.5° for type 3, assuming a helical rise of 4.75 Å. *De novo* 3D initial models were then constructed from 2D class averages representing one whole cross-over of the different filament types using the relion_helix_inimodel2d program (Scheres, 2020). Subsequently, segments were re-extracted without down-sampling in boxes of 256×256 pixels for use in 3D auto-refinements and classifications. Several rounds of refinements were performed, while progressively increasing the resolution of the starting model from 10 Å to 4.5 Å and switching on optimisation of the helical rise and helical twist once β-strands were separated in the starting model. For filaments from seeds of MSA case 1, additional 3D classifications focussed classifications on exterior regions of the filament were used to distinguish the presence of minority polymorphs (with protofilament fold B as described in the main text). Final reconstructions were obtained after Bayesian polishing and CTF refinement, followed by 3D auto-refinement, a 3D classification step without alignment to select the segments contributing to the best classes, a final round of 3D auto-refinement and standard RELION post-processing with a soft solvent mask that extended to 20 % of the box height.

### Atomic modelling

Atomic models of the filaments were built *de novo* in *Coot* (Emsley & Cowtan, 2004) using the maps of the data set for MSA case 2 for type 1 and type 2 filaments with protofilament fold A, and maps of the data set for MSA case 2 for type 1 and type 2 filaments with protofilament fold B. For protofilament fold A, the atomic model with PDB-ID 6UFR of E46K αsynuclein (Boyer et al., 2020) was used as guide. For type 3 filaments, the atomic model with PDB-ID 6PEO (Boyer et al., 2019) of H50Q α-synuclein was used. Models comprising 6 β-sheet rungs were refined in real-space using ISOLDE (Croll, 2018), with interactive flexible molecular dynamics to obtain optimal β-sheet packing chemistry. The resulting models were validated with MolProbity (Chen et al., 2010). Details about the atomic models are described in Table 1.

The schematics in Figure 3 - figure supplement 3e-f and Figure 5 - figure supplement 1b were made with T.Nakane’s atoms2svg.py script, which is publicly available from https://doi.org/10.5281/zenodo.4090924.

---

**Figure 1 – figure supplement 1.**
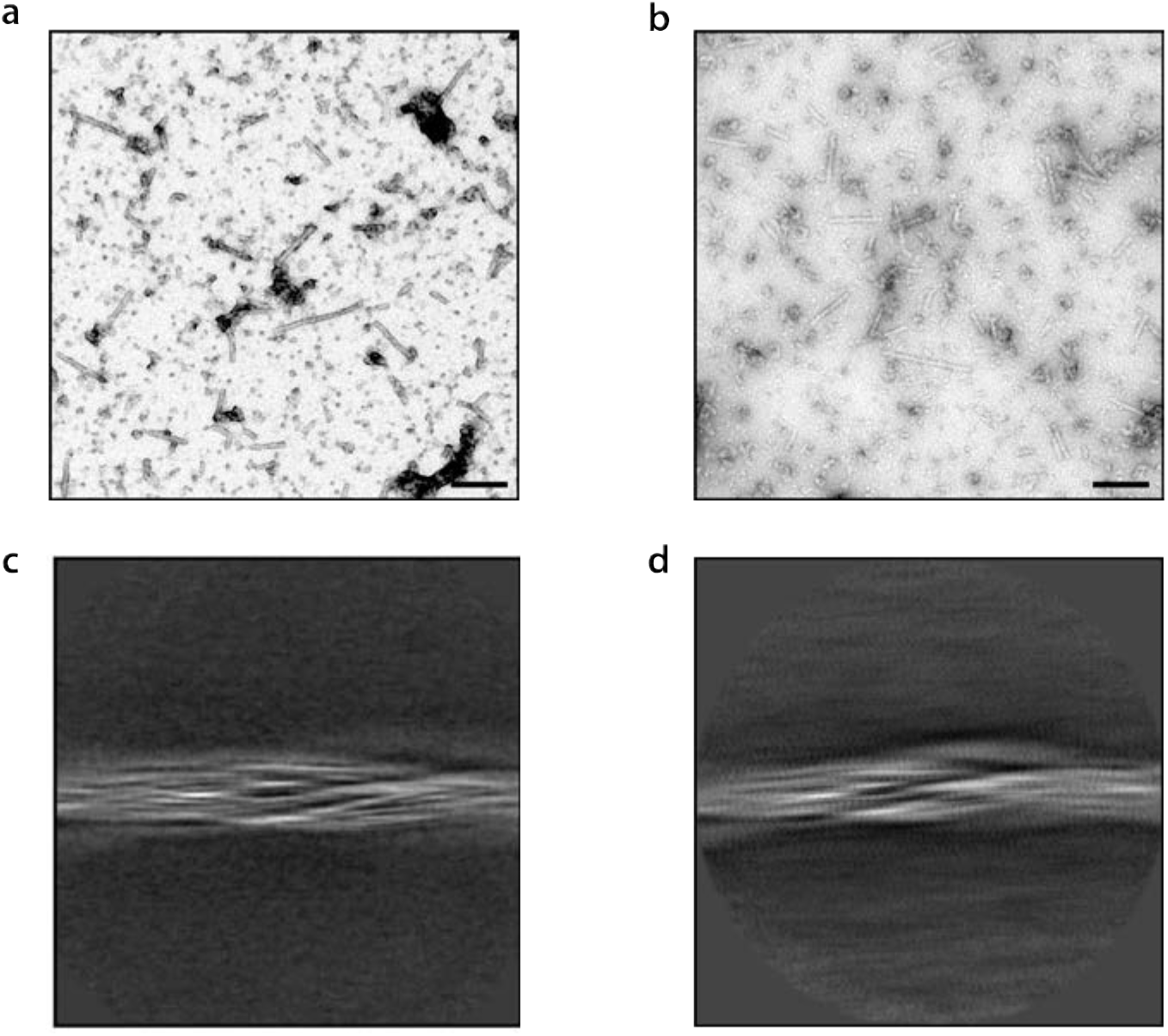
Purified filaments from MSA case 5. **(a)** Sarkosyl insoluble pellet before sonication; and **(b)** after sonication (scale bar = 200 nm). Cryo-EM 2D class averages of MSA case 5 purified filaments before sonication (c) and after sonication (d).

**Figure 2 - figure supplement 1.**
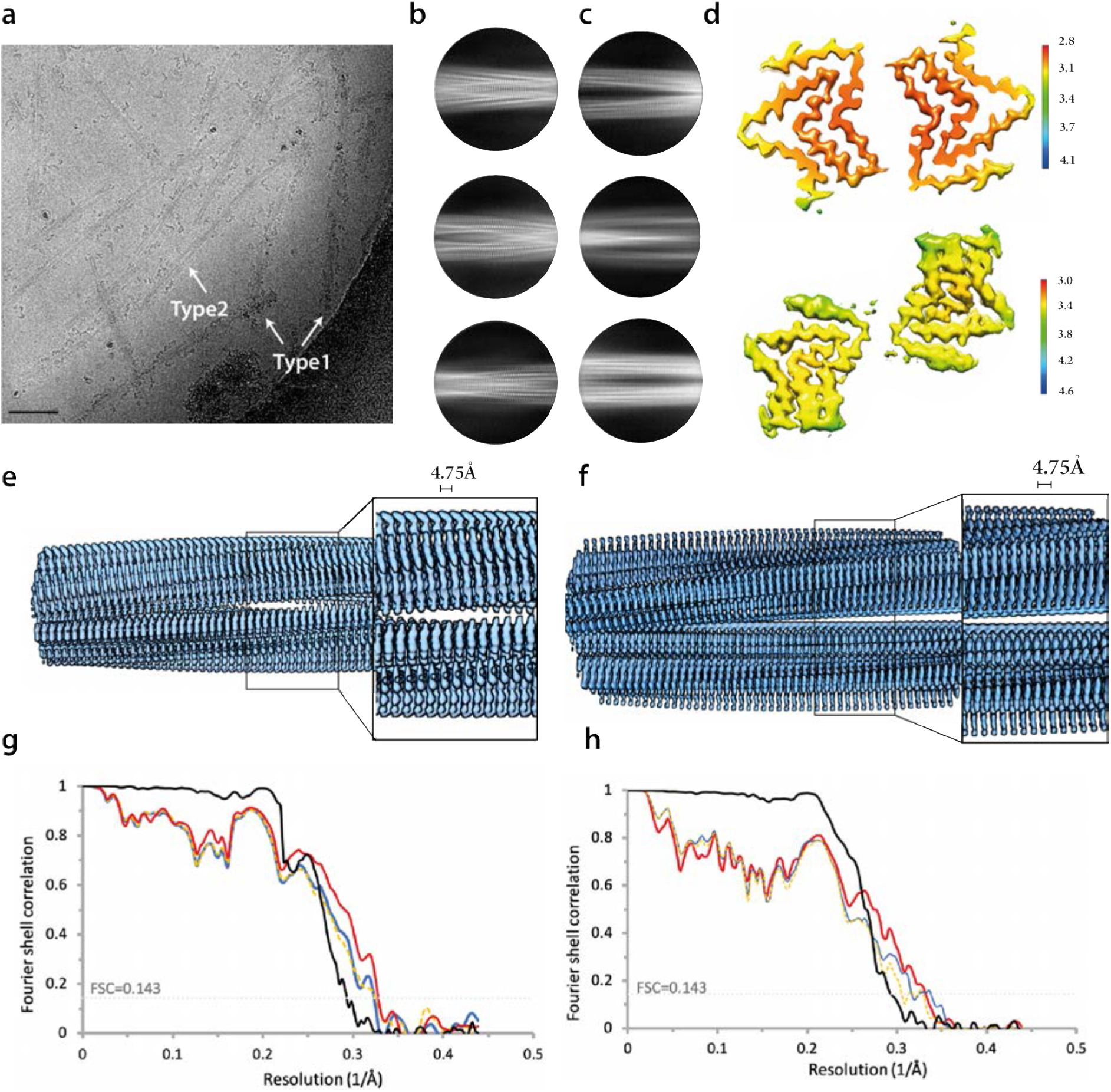
Additional cryo-EM data on type 1 and type 2 filaments with protofilament fold A. **(a)**, Electron micrograph of seeded assemblies using filament preparations from MSA case 2 as the seed. Type 1 and type 2 filaments are indicated with white arrows. Scale bar, 50 nm. 2D class averages of type 1 (left) and type 2 (right) filaments with two protofilaments of fold A in a box spanning 280 Å. **(c)** Local resolution maps for type 1 (top) and type 2 (bottom) filaments, with the legend indicating resolutions in Å. **(d)** Side view of the 3D reconstructions for type 1 (left) and type 2 (right) filaments, showing clear separation of β-strands along the helical axis **(e)** FSC curves for type 1 filaments with two protofilaments of fold A between two independently refined half-maps (black), of the final cryo-EM reconstruction and refined atomic model (red), of the first half map and the atomic model refined against the first half map (blue), and of the atomic model that was refined against the first half-map against the second half-map (yellow dashed). **(f)** As (e), but for type 2 filaments with two protofilaments of fold A.

**Figure 2 - figure supplement 2.**
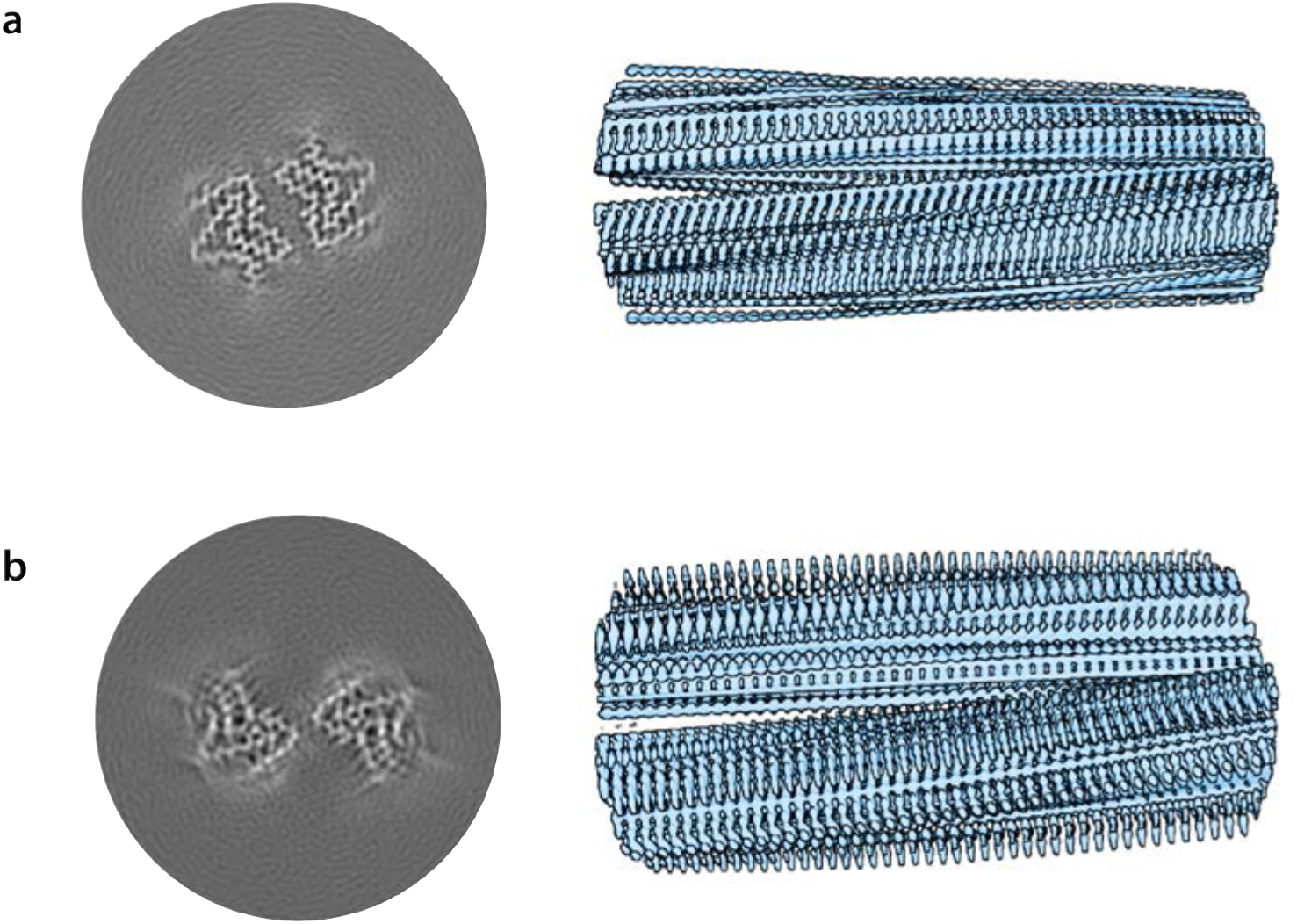
Cryo-EM structures of type 1 and type 2 filaments with protofilament fold A assembled using seeds from MSA case 1. **(a)** Central slice of the 3D map for type 1 filaments. **(b)** Side view of the 3D reconstruction of type 1 filaments. **(c-d)** As (a-b), but for type 2 filaments.

**Figure 2 - figure supplement 3.**
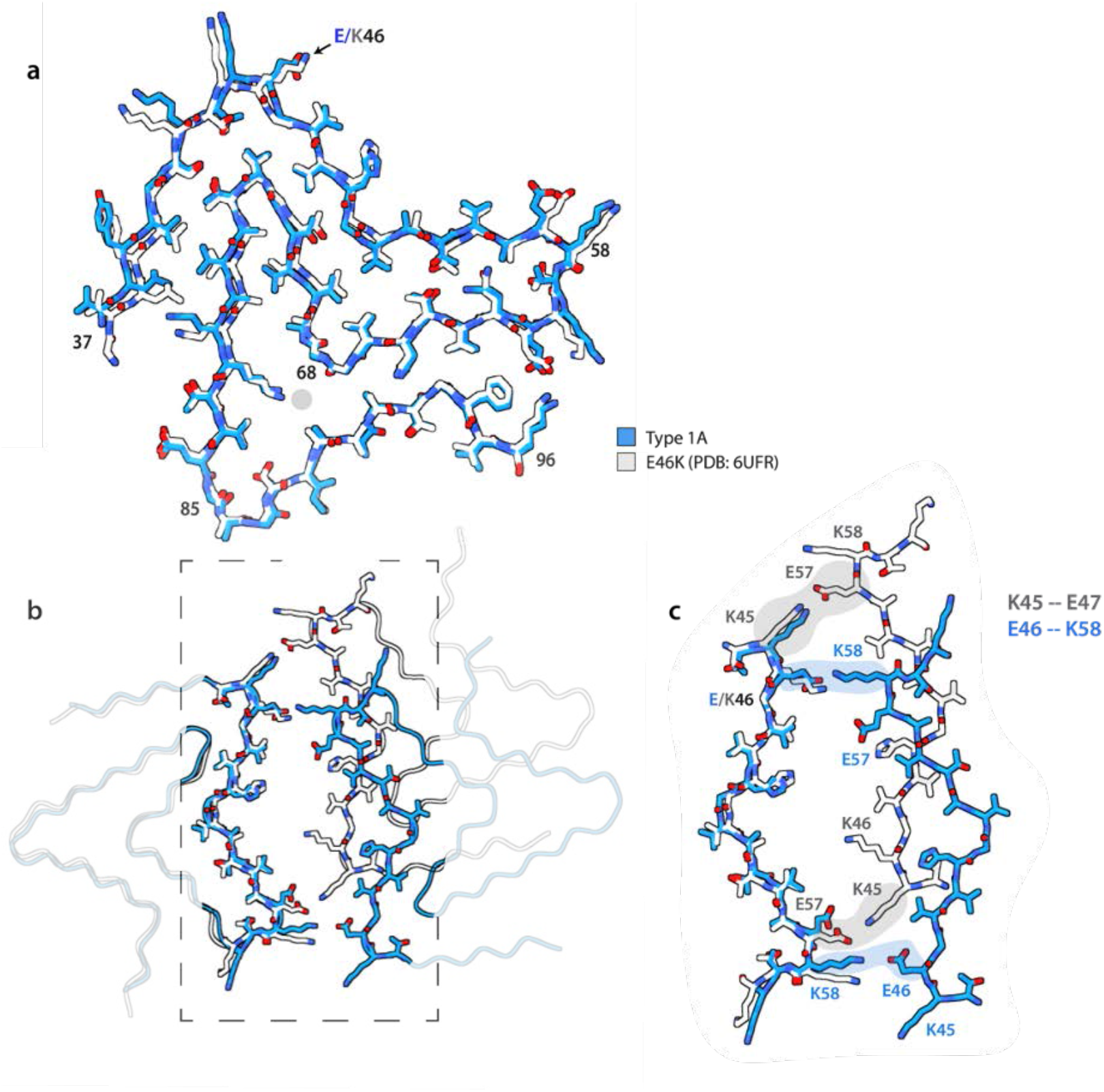
Comparison of protofilament fold A with PDB-entry 6UFR of assembled recombinant E46K α-synuclein. **(a)** Atomic model of protofilament fold A (blue) overlaid with one protofilament from PDB-entry 6UFR (grey). **(b)** Comparison of the interface between two protofilaments with fold A in type 1 filaments and those from PDB entry 6UFR, with the same colour scheme as in (a). (c) Zoomed-in view of the interface, with salt bridges between K45 and E47 in PDB-entry 6UFR and between E46 and K58 in type 1 filaments highlighted in grey and blue, respectively.

**Figure 3 - figure supplement 1.**
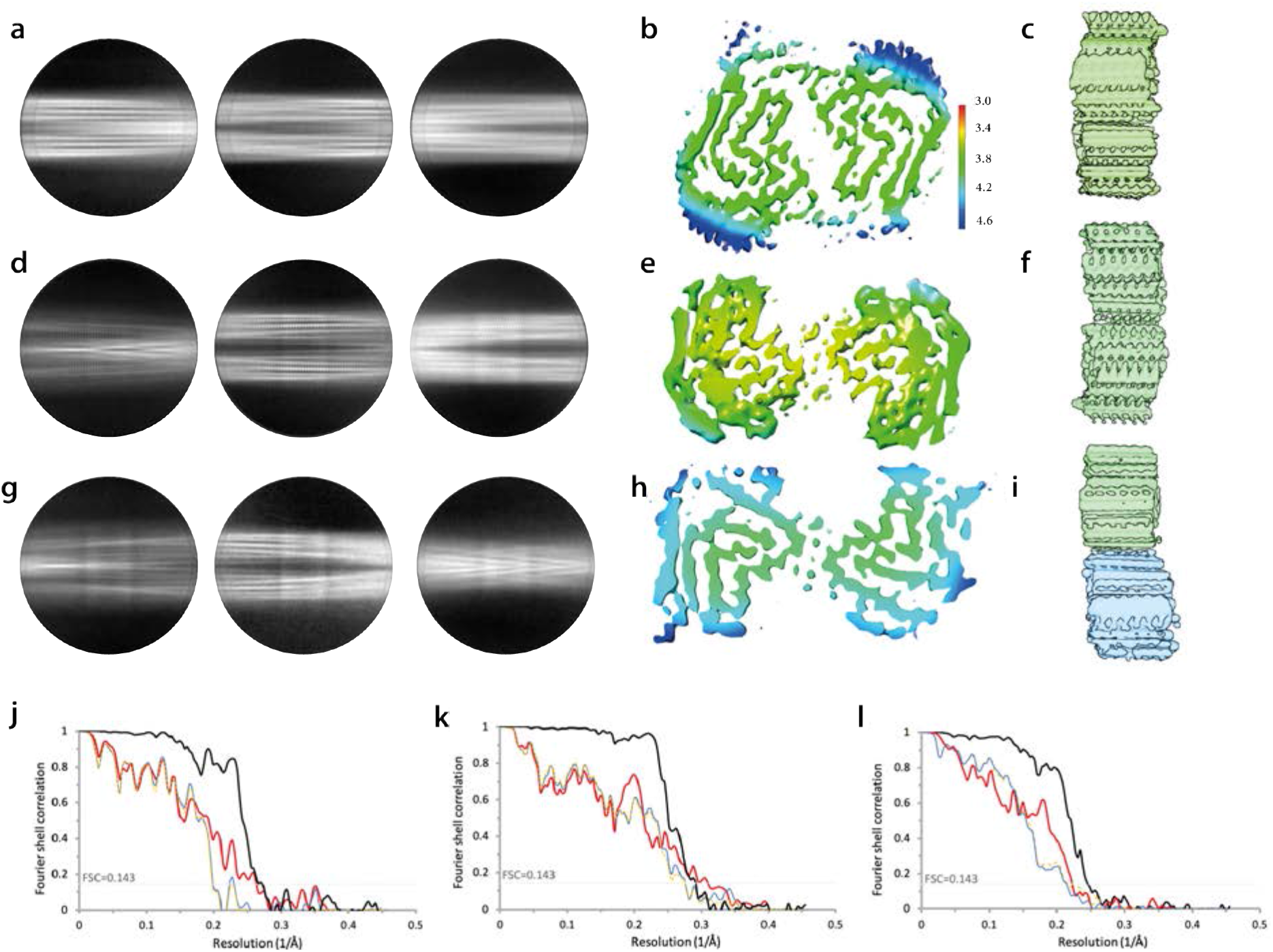
Additional cryo-EM data on type 1 and type 2 filaments with protofilament fold B. **(a)** 2D class averages of type 1 filaments with two protofilaments of fold B **(b)** Local resolution map for type 1 filaments with two protofilaments of fold B with the colour map indicating resolutions in Å. **(c)** Side view of the 3D reconstructions of type 1 filaments with two protofilaments of fold B. **(d-f)** as (a-c) but for type 2 filaments with two protofilaments of fold B. **(g-i)** as (a-c) but for type 2 filaments with one protofilament of fold A and one protofilament of fold B. **(j-l)** Fourier shell correlation curves for type 1 filaments with two protofilaments of fold B (j), type 2 filaments with two protofilaments of fold B (k) and type 2 filaments with one protofilament of fold A and one protofilament of fold B (l). Fourier shell correlation curves are shown between two independently refined half-maps (black) of the final cryo-EM reconstruction and refined atomic model (red), of the first half map and the atomic model refined against the first half map (blue), and of the atomic model that was refined against the first half-map against the second half-map (yellow dashed).

**Figure 3 - figure supplement 2.**
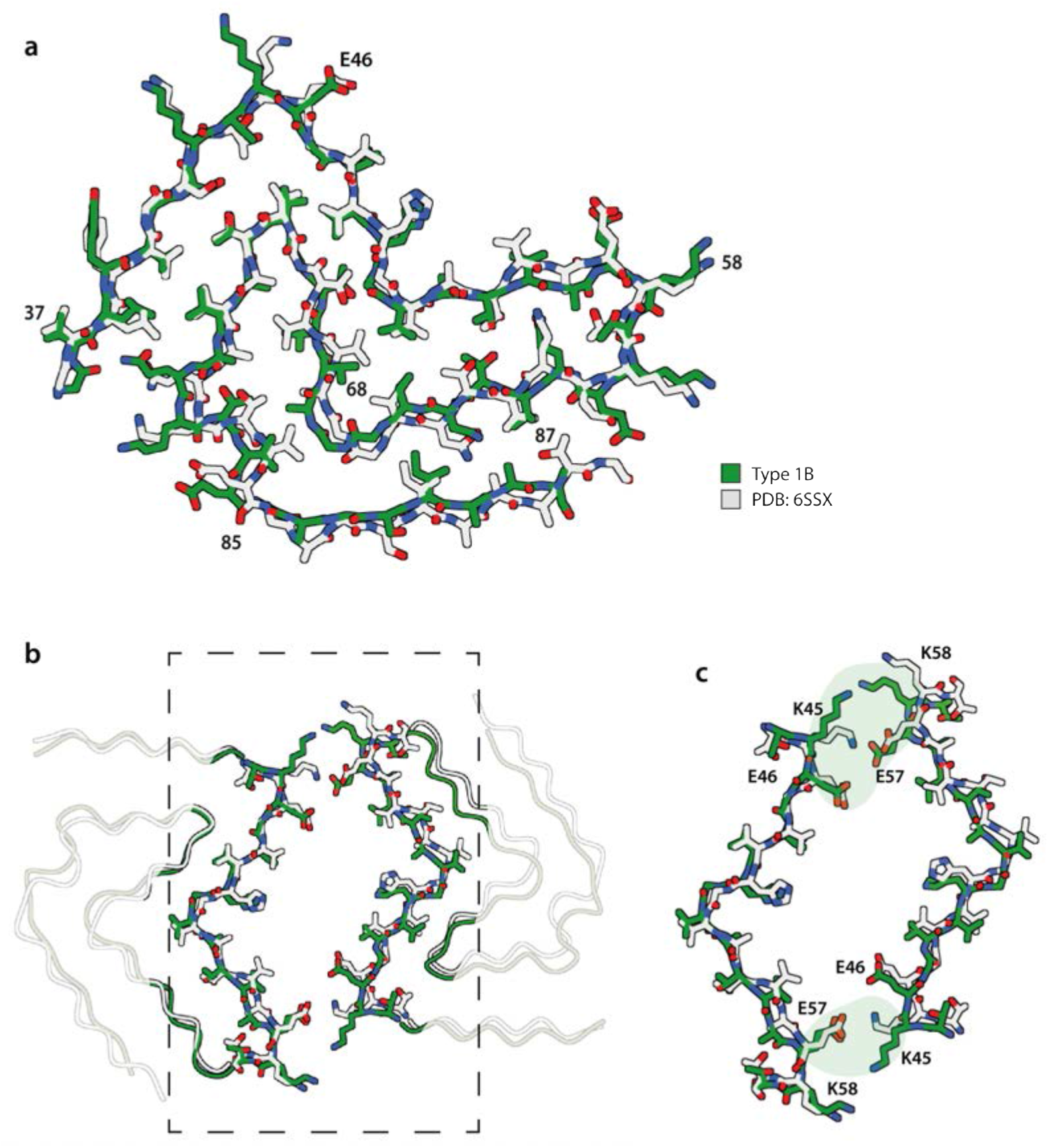
Comparison of protofilament A with PDB-entry 6SSX of recombinant wild-type α-synuclein. **(a)** Atomic model of protofilament fold A (blue) overlaid with one protofilament from PDB-entry 6SSX (grey). **(b)** Comparison of the interface between two protofilaments with fold A in type 1 filaments and those from PDB entry 6UFR, with the same colour scheme as in (a). (c) Zoomed-in view of the interface, with salt bridges between K45 and E47 in PDB-entry 6UPR and between E46 and K58 in type 1 filaments highlighted in grey and blue, respectively.

**Figure 3 - figure supplement 3.**
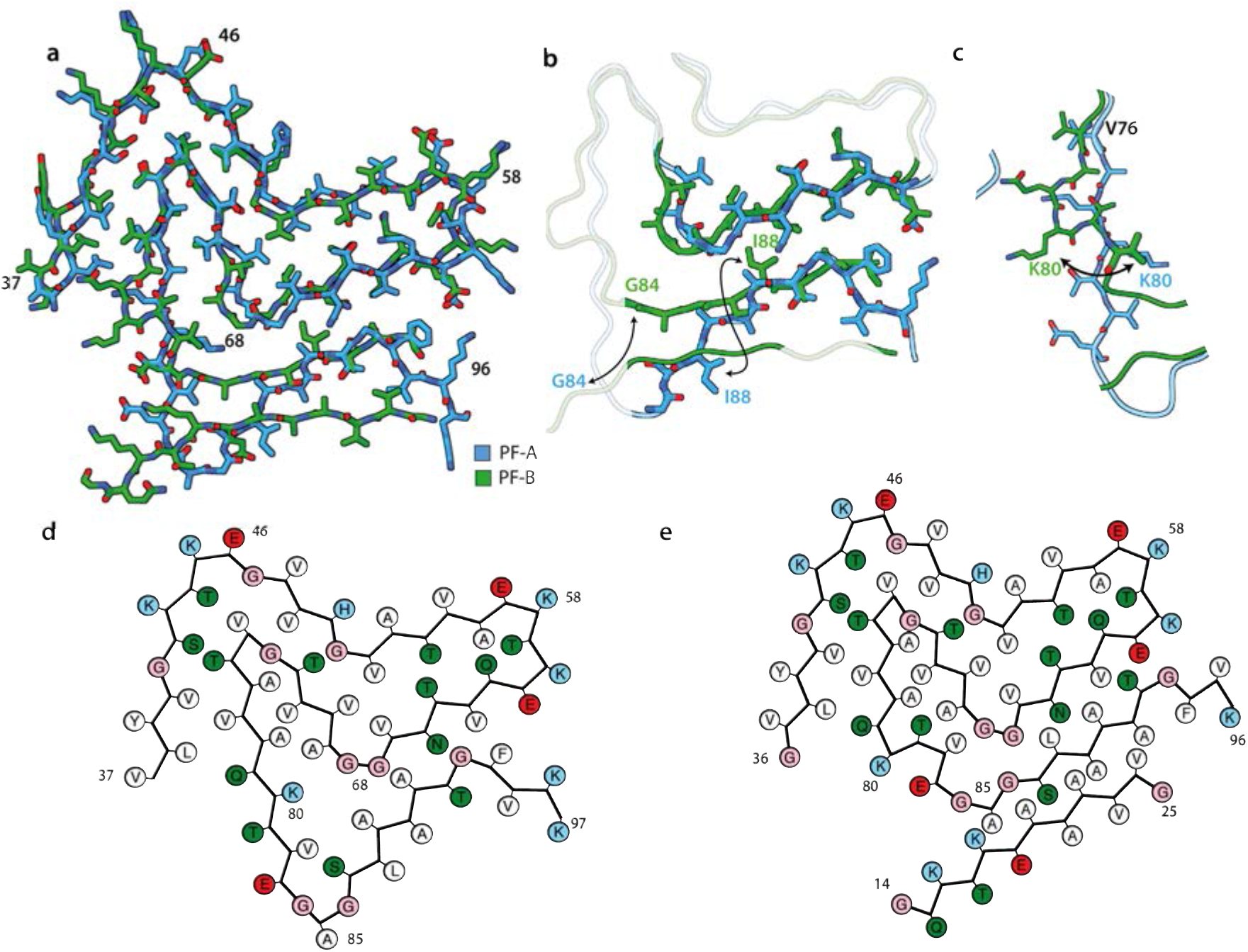
Comparison of protofilament folds A and B. **(a)**Atomic model of protofilament fold A (blue) overlaid with protofilament fold B (green) **(b,c)** As in **(a)**, but showing all-atom representation for different residues. **(d,e)** Schematic representations of protofilament folds A and B. Each amino acid residue is represented with its one-letter code in a circle. Positively charged amino acids are shown in blue, negatively charged ones in red, polar ones in green, hydrophobic ones in white, and glycines in pink.

**Figure 4 - figure supplement 1.**
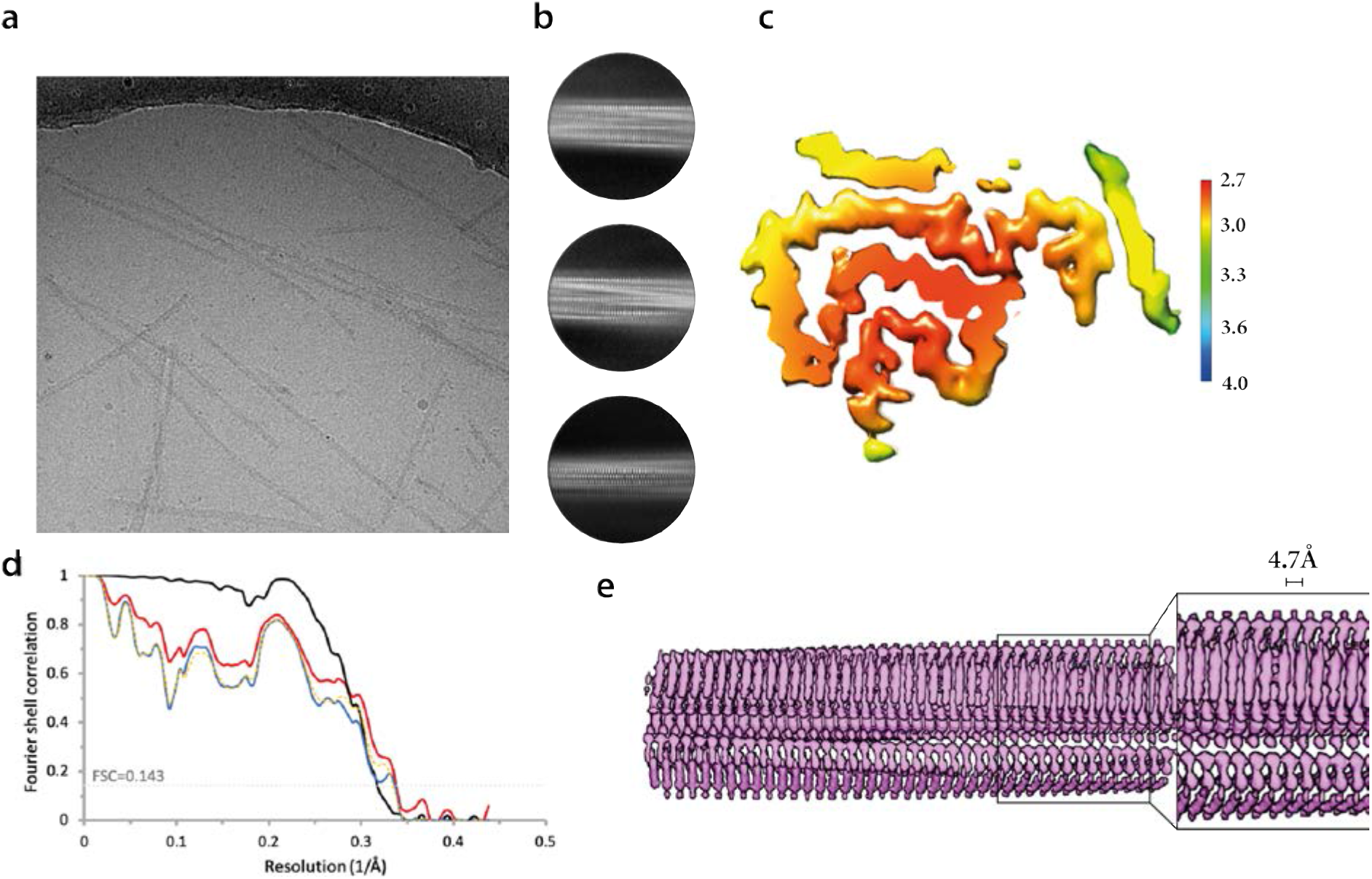
Additional cryo-EM data on type 3 filaments. **(a)** Electron micrograph of case 5. The scale bar indicates 50 nm. **(b)** 2D class averages of type 3 filaments in a box spanning 280 Å. **(c)** Local resolution map, with the colour map indicating resolutions in Å. **(d)**Fourier shell correlation curves between two independently refined half-maps (black), of the final cryo-EM reconstruction and the refined atomic model (red), of the first half map and the atomic model refined against the first half map (blue), and of the atomic model that was refined against the first half-map against the second half-map (yellow dashed). **(e)** Side view of the 3D reconstruction, showing separation of β-strands along the helical axis.

**Figure 4 - figure supplement 2.**
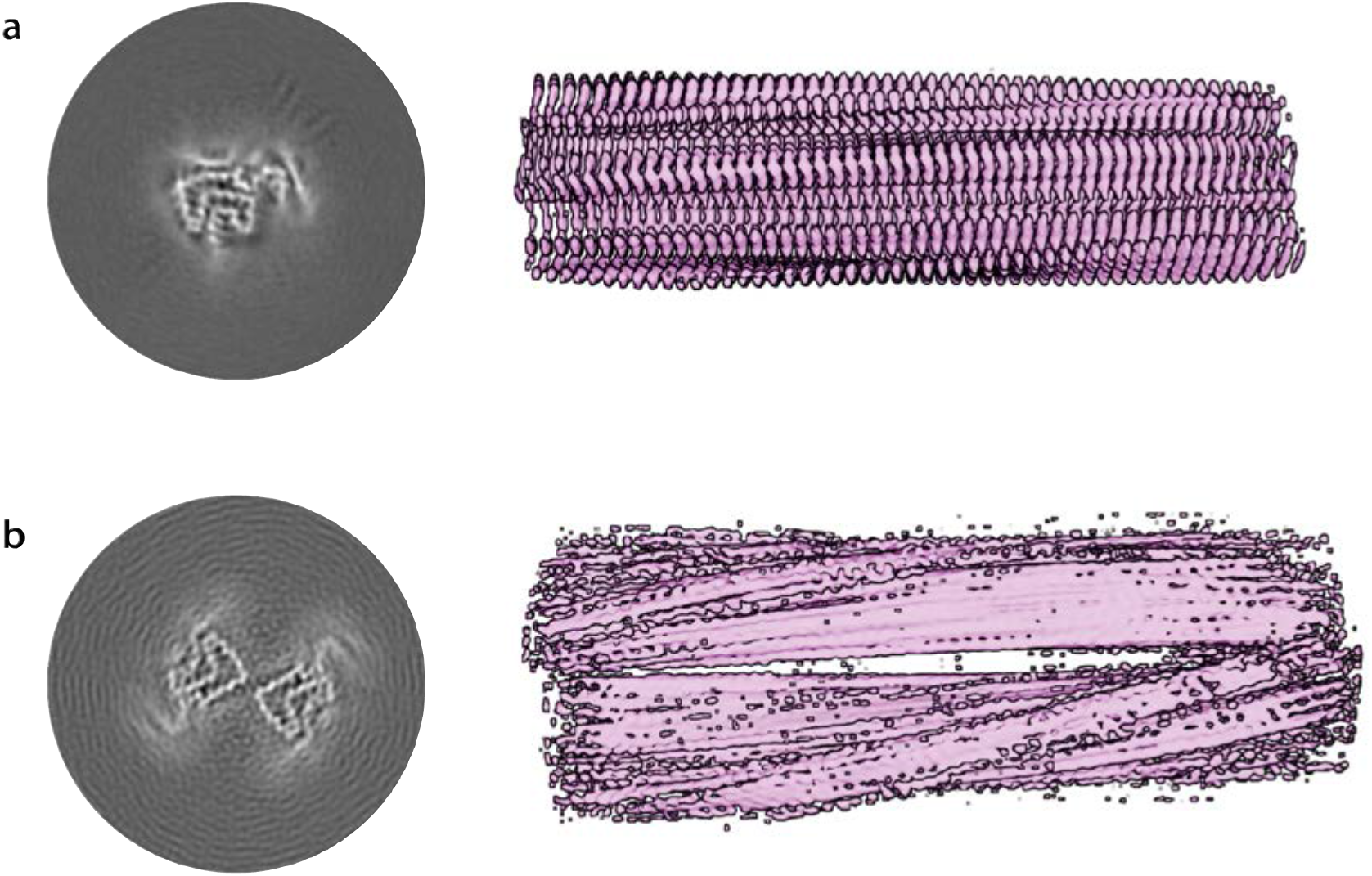
Second-generation type 3 filaments. **(a)** Central slice of the 3D map of the type 3 filaments from the second generation of seeding. Side view of the 3D reconstruction of the same type 3 filaments. **(c-d)** As in (a-b), but for the doublets of type 3 filaments.

**Figure 5 - figure supplement 1.**
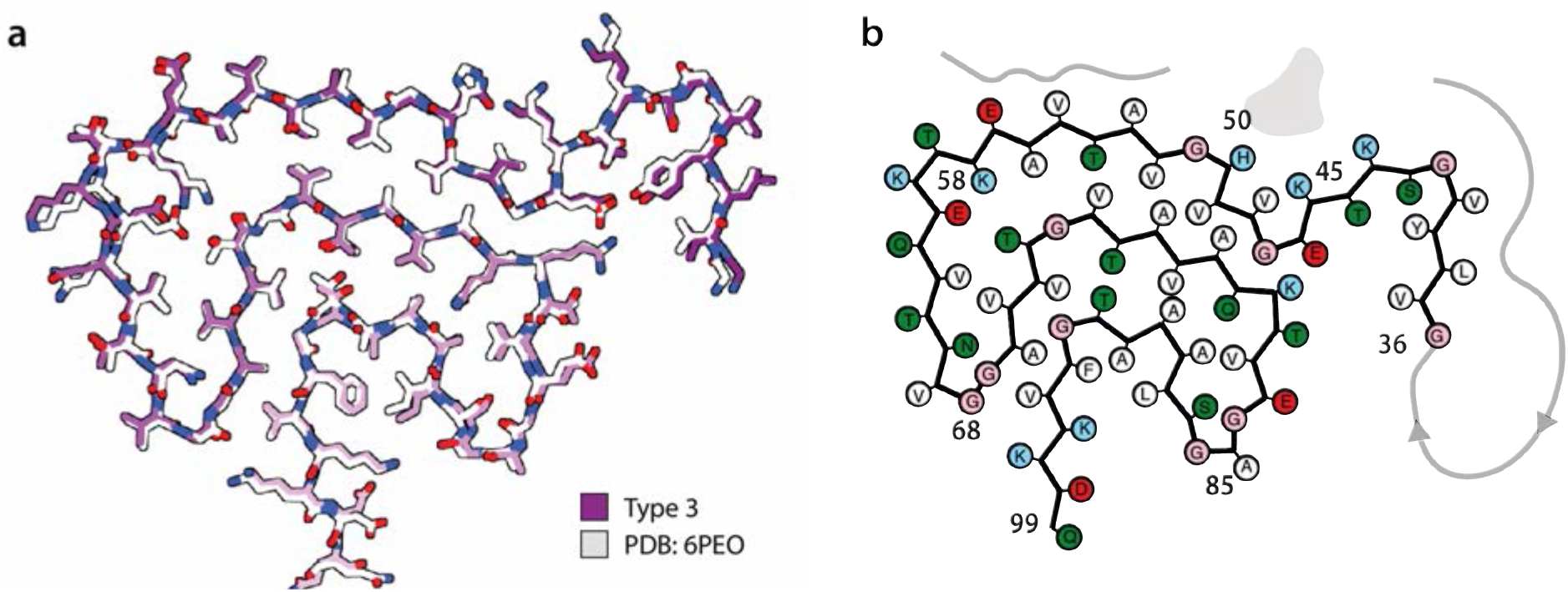
Comparison of type 3 filament with PDB entry 6PEO of assembled recombinant H50Q α-synuclein. **(a)**, All-atom view of the type 3 filament (purple) aligned with PDB-entry 6PEO (grey). **(b)**, Schematic representation of the type 3 filament.

## References

Bassil, F., Fernagut, P.-O., Bezard, E., Pruvost, A., Leste-Lasserre, T., Hoang, Q. Q., Ringe, D., Petsko, G. A., & Meissner, W. G. (2016). Reducing C-terminal truncation mitigates synucleinopathy and neurodegeneration in a transgenic model of multiple system atrophy. Proceedings of the National Academy of Sciences USA, 113(34), 9593–9598. https://doi.org/10.1073/pnas.1609291113

Boyer, D. R., Li, B., Sun, C., Fan, W., Sawaya, M. R., Jiang, L., & Eisenberg, D. S. (2019). Structures of fibrils formed by α-synuclein hereditary disease mutant H50Q reveal new polymorphs. Nature Structural & Molecular Biology 26(11), 1044–1052.

Boyer, D. R., Li, B., Sun, C., Fan, W., Zhou, K., Hughes, M. P., Sawaya, M. R., Jiang, L., & Eisenberg, D. S. (2020). The α-synuclein hereditary mutation E46K unlocks a more stable, pathogenic fibril structure. Proceedings of the National Academy of Sciences USA, 117(7), 3592–3602. https://doi.org/10.1073/pnas.1917914117

Braak, H., & Braak, E. (1991). Neuropathological stageing of Alzheimer-related changes. Acta Neuropathologica, 82(4), 239–259. https://doi.org/10.1007/BF00308809

Braak, Heiko, Tredici, K. D., Rüb, U., de Vos, R. A. I., Jansen Steur, E. N. H., & Braak, E. (2003). Staging of brain pathology related to sporadic Parkinson’s disease. Neurobiology of Aging, 24(2), 197–211. https://doi.org/10.1016/S0197-4580(02)00065-9

Buell, A. K., Galvagnion, C., Gaspar, R., Sparr, E., Vendruscolo, M., Knowles, T. P. J., Linse, S., & Dobson, C. M. (2014). Solution conditions determine the relative importance of nucleation and growth processes in -synuclein aggregation. Proceedings of the National Academy of Sciences USA, 111(21), 7671–7676. https://doi.org/10.1073/pnas.1315346111

Chen, V. B., Arendall, W. B., Headd, J. J., Keedy, D. A., Immormino, R. M., Kapral, G. J., Murray, L. W., Richardson, J. S., & Richardson, D. C. (2010). MolProbity: All-atom structure validation for macromolecular crystallography. Acta Crystallographica. Section D, Biological Crystallography, 66(Pt 1), 12–21. https://doi.org/10.1107/S0907444909042073

Collinge, J., & Clarke, A. R. (2007). A general model of prion strains and their pathogenicity. Science, 318(5852), 930–936. https://doi.org/10.1126/science.1138718

Conway, K. A., Harper, J. D., & Lansbury, P. T. (1998). Accelerated in vitro fibril formation by a mutant α-synuclein linked to early-onset Parkinson disease. Nature Medicine, 4(11), 1318–1320. https://doi.org/10.1038/3311

Croll, T. I. (2018). ISOLDE: A physically realistic environment for model building into low-resolution electron-density maps. Acta Crystallographica Section D: Structural Biology, 74(6), 519–530. https://doi.org/10.1107/S2059798318002425

Crowther, R. A., Jakes, R., Spillantini, M. G., & Goedert, M. (1998). Synthetic filaments assembled from C-terminally truncated α-synuclein. FEBS Letters, 436(3), 309–312. https://doi.org/10.1016/S0014-5793(98)01146-6

Emsley, P., & Cowtan, K. (2004). Coot: Model-building tools for molecular graphics. Acta Crystallographica Section D: Biological Crystallography, 60(12), 2126–2132. https://doi.org/10.1107/S0907444904019158

Falcon, B., Zivanov, J., Zhang, W., Murzin, A. G., Garringer, H. J., Vidal, R., Crowther, R. A., Newell, K. L., Ghetti, B., Goedert, M., & Scheres S., (2019). Novel tau filament fold in chronic traumatic encephalopathy encloses hydrophobic molecules. Nature, 568(7752), 420–423. https://doi.org/10.1038/s41586-019-1026-5

Fitzpatrick, A.W.P., Falcon, B., He, S., Murzin, A.G., Murshudov, G., Garringer, H.J., Crowther, R.A., Ghetti, B., Goedert M. & Scheres, S.H.W. (2017) Cryo-EM structures of tau filaments from Alzheimer’s disease. Nature, 547(7662), 185–190.

Fujiwara, H., Hasegawa, M., Dohmae, N., Kawashima, A., Masliah, E., Goldberg, M. S., Shen, J., Takio, K., & Iwatsubo, T. (2002). Alpha-Synuclein is phosphorylated in synucleinopathy lesions. Nature Cell Biology, 4(2), 160–164. https://doi.org/10.1038/ncb748

Galvagnion, C., Brown, J. W. P., Ouberai, M. M., Flagmeier, P., Vendruscolo, M., Buell, A. K., Sparr, E., & Dobson, C. M. (2016). Chemical properties of lipids strongly affect the kinetics of the membrane-induced aggregation of α-synuclein. Proceedings of the National Academy of Sciences USA, 113(26), 7065–7070. https://doi.org/10.1073/pnas.1601899113

Giasson, B. I., Duda, J. E., Quinn, S. M., Zhang, B., Trojanowski, J. Q., & Lee, V. M.-Y. (2002). Neuronal α-synucleinopathy with severe movement disorder in mice expressing A53T human α-synuclein. Neuron, 34(4), 521–533. https://doi.org/10.1016/S0896-6273(02)00682-7

Goedert, M. (2015). Alzheimer’s and Parkinson’s diseases: The prion concept in relation to assembled Aβ, tau, and α-synuclein. Science, 349(6248), 1255555. https://doi.org/10.1126/science.1255555

Goedert, M., Clavaguera, F., & Tolnay, M. (2010). The propagation of prion-like protein inclusions in neurodegenerative diseases. Trends in Neurosciences, 33(7), 317–325. https://doi.org/10.1016/j.tins.2010.04.003

Goedert, M., Jakes, R., & Spillantini, M.G. (2017). The Synucleinopathies: Twenty years on. Journal of Parkinson’s disease, 7 (S1), 51–69.

Guerrero-Ferreira, R., Kovacik, L., Ni, D., & Stahlberg, H. (2020). New insights on the structure of alpha-synuclein fibrils using cryo-electron microscopy. Current Opinion in Neurobiology, 61, 89–95. https://doi.org/10.1016/j.conb.2020.01.014

Guerrero-Ferreira, R., Taylor, N. M., Arteni, A.-A., Kumari, P., Mona, D., Ringler, P., Britschgi, M., Lauer, M. E., Makky, A., Verasdonck, J., Riek, R., Melki, R., Meier, B. H., Böckmann, A., Bousset, L., & Stahlberg, H. (2019). Two new polymorphic structures of human full-length alpha-synuclein fibrils solved by cryo-electron microscopy. ELife, 8, e48907. https://doi.org/10.7554/eLife.48907

Guerrero-Ferreira, R., Taylor, N. M., Mona, D., Ringler, P., Lauer, M. E., Riek, R., Britschgi, M., & Stahlberg, H. (2018). Cryo-EM structure of alpha-synuclein fibrils. ELife, 7, e36402. https://doi.org/10.7554/eLife.36402

He, S., & Scheres, S. (2017). Helical reconstruction in RELION. Journal of Structural Biology, 198(3), 163–176. https://doi.org/10.1016/j.jsb.2017.02.003

Holec, S. A. M., & Woerman, A. L. (2020). Evidence of distinct α-synuclein strains underlying disease heterogeneity. Acta Neuropathologica. https://doi.org/10.1007/s00401-020-02163-5

Kollmer, M., Close, W., Funk, L., Rasmussen, J., Bsoul, A., Schierhorn, A., Schmidt, M., Sigurdson, C.J., Jucker, M. & Fändrich, M. (2019) Cryo-EM structure and polymorphism of Aβ amyloid fibrils purified from Alzheimer’s brain tissues. Nature Communications, 10: 4760.

Kordower, J. H., Chu, Y., Hauser, R. A., Freeman, T. B., & Olanow, C. W. (2008). Lewy body–like pathology in long-term embryonic nigral transplants in Parkinson’s disease. Nature Medicine, 14(5), 504–506. https://doi.org/10.1038/nm1747

Lavenir, I., Passarella, D., Masuda-Suzukake, M., Curry, A., Holton, J. L., Ghetti, B., & Goedert, M. (2019). Silver staining (Campbell-Switzer) of neuronal α-synuclein assemblies induced by multiple system atrophy and Parkinson’s disease brain extracts in transgenic mice. Acta Neuropathologica Communications, 7(1), 148. https://doi.org/10.1186/s40478-019-0804-5

Li, J., Browning, S., Mahal, S. P., Oelschlegel, A. M., & Weissmann, C. (2010). Darwinian evolution of prions in cell culture. Science (New York, N.Y.), 327(5967), 869–872. https://doi.org/10.1126/science.1183218

Li, J.-Y., Englund, E., Holton, J. L., Soulet, D., Hagell, P., Lees, A. J., Lashley, T., Quinn, N. P., Rehncrona, S., Björklund, A., Widner, H., Revesz, T., Lindvall, O., & Brundin, P. (2008). Lewy bodies in grafted neurons in subjects with Parkinson’s disease suggest host-to-graft disease propagation. Nature Medicine, 14(5), 501–503. https://doi.org/10.1038/nm1746

Luk, K. C., Song, C., O’Brien, P., Stieber, A., Branch, J. R., Brunden, K. R., Trojanowski, J. Q., & Lee, V. M.-Y. (2009). Exogenous α-synuclein fibrils seed the formation of Lewy body-like intracellular inclusions in cultured cells. Proceedings of the National Academy of Sciences USA, 106(47), 20051–20056. https://doi.org/10.1073/pnas.0908005106

Miake, H., Mizusawa, H., Iwatsubo, T., & Hasegawa, M. (2002). Biochemical Characterization of the core structure of α-synuclein filaments. Journal of Biological Chemistry, 277(21), 19213–19219. https://doi.org/10.1074/jbc.M110551200

Morgan, S. A., Lavenir, I., Fan, J., Masuda-Suzukake, M., Passarella, D., DeTure, M. A., Dickson, D. W., Ghetti, B., & Goedert, M. (2020). α-Synuclein filaments from transgenic mouse and human synucleinopathy-containing brains are major seed-competent species. Journal of Biological Chemistry, 295(19), 6652–6664. https://doi.org/10.1074/jbc.RA119.012179

Nalls, M. A., Pankratz, N., Lill, C. M., Do, C. B., Hernandez, D. G., Saad, M., DeStefano, A. L., Kara, E., Bras, J., Sharma, M., Schulte, C., Keller, M. F., Arepalli, S., Letson, C., Edsall, C., Stefansson, H., Liu, X., Pliner, H., Lee, J. H., ... Singleton, A. B. (2014). Large-scale meta-analysis of genome-wide association data identifies six new risk loci for Parkinson’s disease. Nature Genetics, 46(9), 989–993. https://doi.org/10.1038/ng.3043

Ostrerova-Golts, N., Petrucelli, L., Hardy, J., Lee, J. M., Farer, M., & Wolozin, B. (2000). The A53T α-synuclein mutation increases iron-dependent aggregation and toxicity. Journal of Neuroscience, 20(16), 6048–6054. https://doi.org/10.1523/JNEUROSCI.20-16-06048.2000

Papp, M. I., Kahn, J. E., & Lantos, P. L. (1989). Glial cytoplasmic inclusions in the CNS of patients with multiple system atrophy (striatonigral degeneration, olivopontocerebellar atrophy and Shy-Drager syndrome). Journal of Neurological Sciences, 94(1), 79–100. https://doi.org/10.1016/0022-510X(89)90219-0

Peng, C., Gathagan, R. J., Covell, D. J., Medellin, C., Stieber, A., Robinson, J. L., Zhang, B., Pitkin, R. M., Olufemi, M. F., Luk, K. C., Trojanowski, J. Q., & Lee, V. M.-Y. (2018). Cellular milieu imparts distinct pathological α-synuclein strains in α-synucleinopathies. Nature, 557(7706), 558–563. https://doi.org/10.1038/s41586-018-0104-4

Polymeropoulos, M. H., Lavedan, C., Leroy, E., Ide, S. E., Dehejia, A., Dutra, A., Pike, B., Root, H., Rubenstein, J., Boyer, R., Stenroos, E. S., Chandrasekharappa, S., Athanassiadou, A., Papapetropoulos, T., Johnson, W. G., Lazzarini, A. M., Duvoisin, R. C., Iorio, G. D., Golbe, L. I., & Nussbaum, R. L. (1997). Mutation in the α-synuclein gene identified in families with Parkinson’s disease. Science, 276(5321), 2045–2047. https://doi.org/10.1126/science.276.5321.2045

Prusiner, S. (1982). Novel proteinaceous infectious particles cause scrapie. Science, 216(4542), 136–144. https://doi.org/10.1126/science.6801762

Rodriguez, J. A., Ivanova, M. I., Sawaya, M. R., Cascio, D., Reyes, F., Shi, D., Sangwan, S., Guenther, E. L., Johnson, L. M., Zhang, M., Jiang, L., Arbing, M. A., Nannega, B., Hattne, J., Whitelegge, J., Brewster, A. S., Messerschmidt, M., Boutet, S., Sauter, N. K., ... Eisenberg, D. (2015). Structure of the toxic core of α-synuclein from invisible crystals. Nature, 525(7570), 486–490. https://doi.org/10.1038/nature15368

Rohou, A., & Grigorieff, N. (2015). CTFFIND4: Fast and accurate defocus estimation from electron micrographs. Journal of Structural Biology, 192(2), 216–221. https://doi.org/10.1016/j.jsb.2015.08.008

Scheres, S., (2020). Amyloid structure determination in RELION-3.1. Acta Crystallographica Section D: Structural Biology, 76(2), 94–101. https://doi.org/10.1107/S2059798319016577

Scheres, S., Zhang, W., Falcon, B., & Goedert, M. (2020). Cryo-EM structures of tau filaments. Current Opinion in Structural Biology, 64, 17–25. https://doi.org/10.1016/j.sbi.2020.05.011

Schweighauser, M., Shi, Y., Tarutani, A., Kametani, F., Murzin, A. G., Ghetti, B., Matsubara, T., Tomita, T., Ando, T., Hasegawa, K., Murayama, S., Yoshida, M., Hasegawa, M., Scheres, S., & Goedert, M. (2020). Structures of α-synuclein filaments from multiple system atrophy. Nature, 585(7825), 464–469. https://doi.org/10.1038/s41586-020-2317-6

Serpell, L. C., Berriman, J., Jakes, R., Goedert, M., & Crowther, R. A. (2000). Fiber diffraction of synthetic alpha-synuclein filaments shows amyloid-like cross-beta conformation. Proceedings of the National Academy of Sciences USA, 97(9), 4897–4902. https://doi.org/10.1073/pnas.97.9.4897

Shahnawaz, M., Mukherjee, A., Pritzkow, S., Mendez, N., Rabadia, P., Liu, X., Hu, B., Schmeichel, A., Singer, W., Wu, G., Tsai, A.-L., Shirani, H., Nilsson, K. P. R., Low, P. A., & Soto, C. (2020). Discriminating α-synuclein strains in Parkinson’s disease and multiple system atrophy. Nature, 578(7794), 273–277. https://doi.org/10.1038/s41586-020-1984-7

Singleton, A. B., Farrer, M., Johnson, J., Singleton, A., Hague, S., Kachergus, J., Hulihan, M., Peuralinna, T., Dutra, A., Nussbaum, R., Lincoln, S., Crawley, A., Hanson, M., Maraganore, D., Adler, C., Cookson, M. R., Muenter, M., Baptista, M., Miller, D., ... Gwinn-Hardy, K. (2003). α-Synuclein locus triplication causes Parkinson’s disease. Science, 302(5646), 841–841. https://doi.org/10.1126/science.1090278

Sorrentino, Z. A., & Giasson, B. I. (2020). The emerging role of α-synuclein truncation in aggregation and disease. Journal of Biological Chemistry, 295(30), 10224–10244. https://doi.org/10.1074/jbc.REV120.011743

Spillantini, M. G., Crowther, R. A., Jakes, R., Hasegawa, M., & Goedert, M. (1998). α-Synuclein in filamentous inclusions of Lewy bodies from Parkinson’s disease and dementia with Lewy bodies. Proceedings of the National Academy of Sciences USA, 95(11), 6469–6473.

Strohäker, T., Jung, B. C., Liou, S.-H., Fernandez, C. O., Riedel, D., Becker, S., Halliday, G. M., Bennati, M., Kim, W. S., Lee, S.-J., & Zweckstetter, M. (2019). Structural heterogeneity of α-synuclein fibrils amplified from patient brain extracts. Nature Communications, 10, 5535 https://doi.org/10.1038/s41467-019-13564-w

Tarutani, A., Arai, T., Murayama, S., Hisanaga, S.-I., & Hasegawa, M. (2018). Potent prion-like behaviors of pathogenic α-synuclein and evaluation of inactivation methods. Acta Neuropathologica Communications, 6(1), 29. https://doi.org/10.1186/s40478-018-0532-2

Tarutani, A., Suzuki, G., Shimozawa, A., Nonaka, T., Akiyama, H., Hisanaga, S.-I., & Hasegawa, M. (2016). The effect of fragmented pathogenic α-synuclein seeds on prion-like propagation. Journal of Biological Chemistry, 291(36), 18675–18688. https://doi.org/10.1074/jbc.M116.734707

Tuttle, M. D., Comellas, G., Nieuwkoop, A. J., Covell, D. J., Berthold, D. A., Kloepper, K. D., Courtney, J. M., Kim, J. K., Barclay, A. M., Kendall, A., Wan, W., Stubbs, G., Schwieters, C. D., Lee, V. M. Y., George, J. M., & Rienstra, C. M. (2016). Solid-state NMR structure of a pathogenic fibril of full-length human α-synuclein. Nature Structural & Molecular Biology, 23(5), 409–415. https://doi.org/10.1038/nsmb.3194

Vilar, M., Chou, H.-T., Lührs, T., Maji, S. K., Riek-Loher, D., Verel, R., Manning, G., Stahlberg, H., & Riek, R. (2008). The fold of α-synuclein fibrils. Proceedings of the National Academy of Sciences USA, 105(25), 8637–8642. https://doi.org/10.1073/pnas.0712179105

Watts, J. C., Giles, K., Oehler, A., Middleton, L., Dexter, D. T., Gentleman, S. M., DeArmond, S. J., & Prusiner, S. B. (2013). Transmission of multiple system atrophy prions to transgenic mice. Proceedings of the National Academy of Sciences USA, 110(48), 19555–19560. https://doi.org/10.1073/pnas.1318268110

Woerman, A. L., Kazmi, S. A., Patel, S., Aoyagi, A., Oehler, A., Widjaja, K., Mordes, D. A., Olson, S. H., & Prusiner, S. B. (2018). Familial Parkinson’s point mutation abolishes multiple system atrophy prion replication. Proceedings of the National Academy of Sciences USA, 115(2), 409–414. https://doi.org/10.1073/pnas.1719369115

Woerman, A. L., Stöhr, J., Aoyagi, A., Rampersaud, R., Krejciova, Z., Watts, J. C., Ohyama, T., Patel, S., Widjaja, K., Oehler, A., Sanders, D. W., Diamond, M. I., Seeley, W. W., Middleton, L. T., Gentleman, S. M., Mordes, D. A., Südhof, T. C., Giles, K., & Prusiner, S. B. (2015). Propagation of prions causing synucleinopathies in cultured cells. Proceedings of the National Academy of Sciences USA, 112, E4949–E4958 https://doi.org/10.1073/pnas.1513426112

Xue, C., Lin, T. Y., Chang, D., & Guo, Z. (2017). Thioflavin T as an amyloid dye: Fibril quantification, optimal concentration and effect on aggregation. Royal Society Open Science, 4(1). https://doi.org/10.1098/rsos.160696

Yonetani, M., Nonaka, T., Masuda, M., Inukai, Y., Oikawa, T., Hisanaga, S.-I., & Hasegawa, M. (2009) Conversion of wild-type α-synuclein into mutant-type fibrils and its propagation in the presence of A30P mutant. Journal of Biological Chemistry, 287(12), 7940–7950.

Zhang, W., Falcon, B., Murzin, A. G., Fan, J., Crowther, R. A., Goedert, M., & Scheres, S. H. (2019). Heparin-induced tau filaments are polymorphic and differ from those in Alzheimer’s and Pick’s diseases. ELife, 8, e43584. https://doi.org/10.7554/eLife.43584

Zhao, K., Lim, Y.-J., Liu, Z., Long, H., Sun, Y., Hu, J.-J., Zhao, C., Tao, Y., Zhang, X., Li, D., Li, Y.-M., & Liu, C. (2020). Parkinson’s disease-related phosphorylation at Tyr39 rearranges α-synuclein amyloid fibril structure revealed by cryo-EM. Proceedings of the National Academy of Sciences USA, 117(33), 20305–20315.

Zivanov, J., Nakane, T., Forsberg, B. O., Kimanius, D., Hagen, W. J., Lindahl, E., & Scheres S., (2018). New tools for automated high-resolution cryo-EM structure determination in RELION-3. ELife, 7, e42166. https://doi.org/10.7554/eLife.42166

Zivanov, J., Nakane, T., & Scheres, S., (2020). Estimation of high-order aberrations and anisotropic magnification from cryo-EM data sets in RELION-3.1. IUCrJ, 7(Pt 2), 253–267. https://doi.org/10.1107/S2052252520000081

